# Competing signaling pathways controls electrotaxis

**DOI:** 10.1101/2025.01.10.631288

**Authors:** S. Kulkarni, F. Tebar, C. Rentero, M. Zhao, P. Sáez

## Abstract

Understanding how cells migrate following exogenous cues is one of the most fundamental questions for biology, medicine, and bioengineering. Growing evidence shows that electrotaxis, the directed cell migration toward electric potential gradients, represents a precise and programmable method to control cell migration. Most data suggest that the polarization of membrane components and the following downstream signaling are central to electrotaxis. Unfortunately, how these multiple mechanisms coordinate with the motile machinery of the cell to respond to an electric field is still poorly understood. Here, we develop a mechanistic model that explains and recapitulate electrotaxis across different cell types. Using the zebrafish proteome, we identify membrane proteins directly related to migration signaling pathways that polarize anodally and cathodally. Further, we show that simultaneous and asymmetric distribution of charged membrane receptors towards the anode and the cathode establishes multiple cooperative and competing stimuli for downstream signaling pathways. The resulting polarization of signals controls the actomyosin network dynamics, directing the anodal and cathodal migration of the cell. Our theoretical framework rationalizes the physical processes that determine electrotaxis across cell types and provides a physical framework to test multiple electrotactic pathways. These results together show us how to control cell migration to, e.g., enhance, cancel, or switch directed cell migration, which opens up new avenues not only to promote tissue regeneration or arrest tumor progression but also to design better biomimetic-engineered tissue constructs.

## Introduction

Cell migration determines some of the most fundamental processes of life, such as embryonic development, wound healing, or tumor invasion, among many others Friedl and Gilmour (2009); Mayor and Etienne-Manneville (2016); Van Helvert et al. (2018). Cells migrate guided by exogenous signals in-vivo and in-vitro. For decades, there have been tremendous efforts to understand how cells organize themselves and with others to follow stimuli in the form of chemical Van Haastert and Devreotes (2004) and mechanical cues Sunyer and Trepat (2020); Shellard and Mayor (2021). There is also an increasing interest in the life sciences and bioengineering in controlling cell migration through external cues because it may allow us to, e.g., arrest tumor cell invasion or promote tissue regeneration Butcher et al. (2009); Friedl and Gilmour (2009); Levin (2009). It may also allow us to precisely design the cellular structure in tissue constructs to achieve remarkable biomimetic features Langer and Tirrell (2004); Wegst et al. (2014).

Electrotaxis, the migration of cells under the influence of electric fields (EFs), represents a powerful and programmable form of guiding cell migration McCaig et al. (2005); Cortese et al. (2014); Zajdel et al. (2020). Numerous experimental observations are consistent with the hypothesis of electrotaxis. Indeed, electrotaxis is ubiquitous across different cell types, including cancer cells Yan et al. (2009), neurons Patel and Poo (1982) or leukocytes Zhao et al. (2006), and tissues Cohen et al. (2014); Zhang et al. (2022). It also appears in different physiological conditions, e.g., during embryonic development Hotary and Robinson (1994) or wound healing Zhao (2009). Most studies have shown that cells migrate toward the cathode. However, we can also find cell types, or cells under specific conditions, that migrate toward the anode. Corneal Epithelial Cells (CECs) migrate to the cathode under the influence of an EF of 150 mV/mm in magnitude, which has been attributed to a cathodal distribution of Epidermal Growth Factor Receptor (EGFR) and membrane lipids Zhao et al. (1999). This behavior was also found in Mammalian Epithelial Cells (MECs) under the same EF strength Zhao et al. (2002b), with EGFR polarization and migration toward the cathode. Embryonic and adult neural progenitor cells show directed migration to the cathode in an EF of 500 mV/mm, which was reduced when PI3K/Atk was inhibitedMeng et al. (2011). Human Telomerase-immortalized Corneal Epithelial (hTCEpi) cells also undergo cathodal electrotaxis in the presence of an EF of 101.2 mV/mm Zhao et al. (2014), as well as in fish keratocytes Sun et al. (2018) and macrophages Sun et al. (2019) in the presence of an EF of 400 mV/mm. Other studies showed that Chinese Hamster Ovary (CHO) cells migrate to the cathode or anode in an electric field of 300 mV/mm depending on the presence of integrins Zhu et al. (2019). EFs of around 75-100 mV/mm cause anodal migration of endothelial cells with an anodal polarization of Vascular Endothelial Growth Factor (VEGF) receptors, PI3K-Akt, and Rho signaling Zhao et al. (2004). The same consistency in the migration and polarization of EGFR toward the anode was found in breast cancer cells Pu et al. (2007); Wu et al. (2013). Bone marrow mesenchymal stem cells and macrophages also show anodal migration Zhao et al. (2011); Sun et al. (2019). Lung cancer cell lines, CL1–5 (highly metastatic) and CL1–0 (weakly metastatic) have shown anodal and weak electrotaxis, respectively Huang et al. (2009). However, it was later demonstrated that CL1-5 cells accumulate EGFR and filopodia on the cathodal side Wang et al. (2011) in an EGFR-independent migration mode Tsai et al. (2013). On cell aggregates, glioblastoma and medulloblastoma migrate cathodally and anodally, respectively, in the same culture and stimulation conditions Lyon et al. (2019).

Unfortunately, how cells follow the EF stimulus is not well understood, and, therefore, important answers to the following questions remain elusive. For example, why do some cells migrate toward the anode while others do toward the cathode? What are the cell sensors that trigger electrotaxis? Are there universal mechanisms that explain electrotaxis across cell types? To answer these questions, we must understand three fundamental layers in the electrotaxis process: first, how an EF is sensed by the cell; second, how that signal is transferred intracellularly to finally control the migration machinery of the cell. We show a schematic representation of the main mechanisms involved in electrotaxis in Fig. 1.

**Figure 1:**
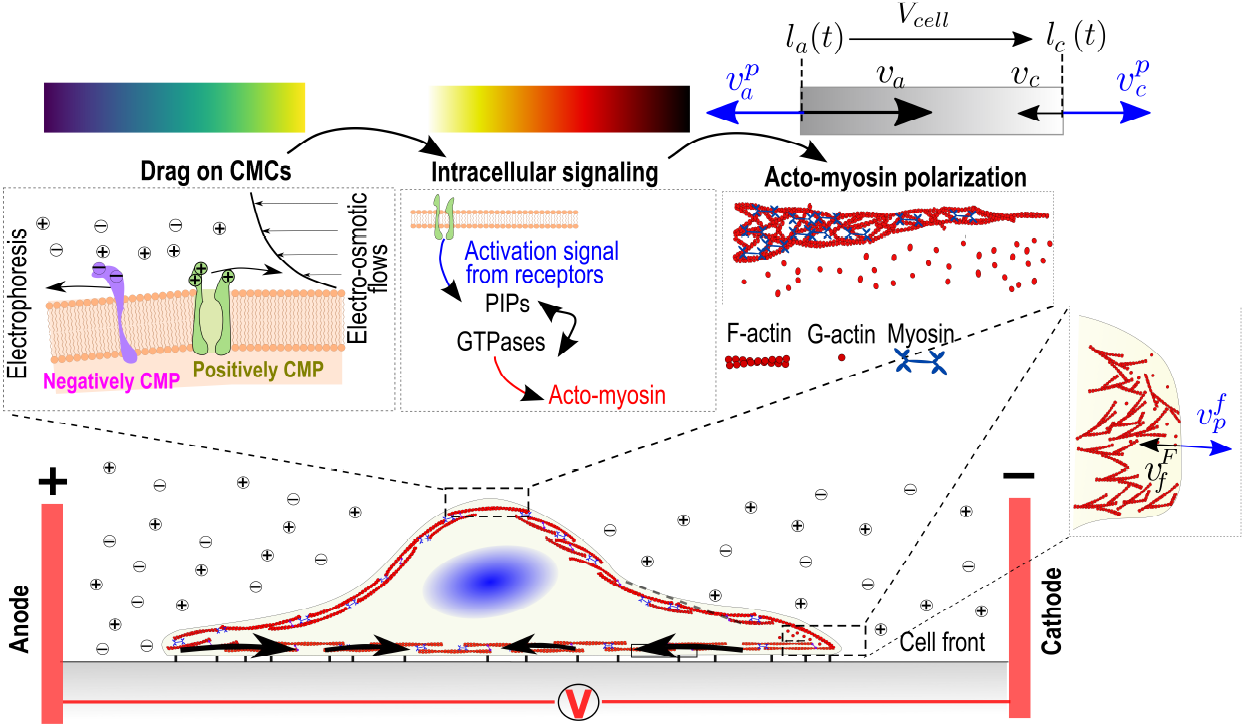
Sketch of the main processes involved in electrotaxis. Two competing forces polarize charged cell membrane components (left). First, an EF exerts forces on the net electric charge of the cell membrane, pulling it in a certain direction (electrophoresis). The generation of an electroosmotic flow by the EF that transports soluble ions and fluid outside the cell (electroosmosis) also generates drag forces on the membrane proteins. The polarization of membrane components induces intracellular signals (upper center) involved in the cell polarization of the actomyosin network that controls the direction of migration. Our 1D electrotaxis model couples CMPs polarization, intracellular signals, and active models (see *Materials and Methods* for details). The retrograde flow, *υ*^*F*^, (black arrows) moves from the cathodal and anodal cell front inwards (subindexes c and a in the variables, respectively). The polymerization velocity (*υ*_*p*_, blue arrow) points outwards from the cell membrane.

When an EF is applied to a cell, only cell components with an extracellular domain can sense it because the EF cannot enter the electrically insulated cytoplasm Mossop et al. (2004). Therefore, cell membrane proteins are likely candidates for sensing an externally applied EF. Among several hypotheses, theoretical and experimental evidence has shown that electrotaxis is consistent with the electromotility model Zhao et al. (2002a); Allen et al. (2013); McCaig et al. (2009), where the electromotility of Charged Membrane Proteins (CMPs) with extracellular domains polarize under the effect of an EF using electrophoresis and electroosmotic forces (see Allen et al. (2013); Sarkar et al. (2019) and references therein). VEGF/EGF receptors Zhao et al. (2002a), integrins Zhu et al. (2019), Receptor tyrosine kinases (RTKs) Jékely et al. (2005) and G protein-coupled receptors (GPCRs) have been shown to polarize under EFs. Many of these proteins then trigger cell polarization through putative downstream signaling adaptation processes Fukata et al. (2003); MacHacek et al. (2009); Ridley (2015). Some of these CMPs can also cluster in lipid rafts, creating complex compounds of molecules that also electro-migrate under an EF Zhao et al. (2002a); Lin et al. (2017).

Mechanistically, an EF, as any other tactic cue, must eventually control the physical forces that dictate cell migration. Cells migrate because of asymmetric intracellular forces that direct cells in specific directions. The forces involved in cell migration can be reduced to three. First, forces that protrude the cell membrane forward due to a continuous polymerization of actin filaments Footer et al. (2007); Schreiber et al. (2010); Wong et al. (2014). The extension of actin filaments, particularly at their fast-growing “barbed ends,” is tightly regulated by various factors involved in nucleation, capping, and depolymerization of actin filaments Small et al. (2002); Pollard et al. (2000). Second, contractile forces generated by myosin motors induce an inward retrograde flow in the actomyosin network Cramer (1997); Pantaloni et al. (2000); Pollard and Cooper (2009). And, third, adhesion forces that are transmitted from the internal actomyosin network to the extracellular space through molecules in the adhesion complexes Parsons et al. (2010); Vicente-Manzanares et al. (2009). Stronger adhesions mean higher effective friction in the retrograde flow which, consequently, slows it down while weak adhesions allow for faster retrograde flows Chan and Odde (2008); Elosegui-Artola et al. (2016); Venturini and Sáez (2023). The polarization and balance of these three forces ultimately determine the strength and direction of cell migration.

Connecting the initial polarization of CMPs in response to exogenous EF and the final polarization of any of those three motile forces, there are intermediate signaling layers. Specifically, small GTPases of the rho family, phosphoinositide kinases, and phosphoinositides phosphate (PIPs) have fundamental roles in cell polarization and migration Ridley et al. (2003); Devreotes and Horwitz (2015). Intricate feedback loops between Rho-GTPases, PIPs, phosphoinositides, and their activating stimuli determine what will become the front and back of the cell and, eventually, the cell migration direction. Understanding the coordination of these pathways in space and time is a fundamental goal for studying electrotaxis and cell migration in general. Further details and modeling options will be described in the following sections.

In short, there is direct experimental evidence of electrotaxis across multiple cell types. However, what specific sensors at the cell membrane trigger electrotaxis, how these CMPs first redistribute, and how they polarize the downstream signals that control the motility forces of the cell to, eventually, establish either an anodal or cathodal migration remains poorly understood. Here, we explore a mechanistic computational model to answer these questions by integrating models of electromigration of CMPs, signaling pathways, and cell migration which, as far as we know, has never been described before. This approach provides a complete, efficient, and adaptable way of analyzing all the elements that cooperate toward electrotaxis. To make this tool available for guiding future experimental research and further modeling efforts, we present the computational model in an online platform uploaded in [the MATLAB Central File Exchange](https://es.mathworks.com/matlabcentral/fileexchange/166511-electrotaxis_1d).

## Results

### Polarization of CMPs and intracellular signals explain electrotaxis

To analyze electrotaxis through our computational model, we initially define an unpolarized state of the cell, i.e. density of CMPs, intracellular signals, actin, and myosin are homogeneously distributed along the cell. The cell expands symmetrically as previously described Betorz et al. (2023) until we stimulate the cell with a physiological EF of 120 mV/mm at t=180s (Fig. 2). We first analyze the polarization of CMPs as the first responders to the externally applied EF. We use an electromigration model to evaluate the redistribution of CMPs. We use values of the *ζ*-potentials of the membrane surface in fibroblasts and mesenchymal stem cells, which are ≈ -60mV Lin et al. (2017). In terms of the CMPs, we first focus on EGFR because it has been shown repetitively as one of the main extracellular components involved in electrotaxis Zhao et al. (1999, 2002b); Wang et al. (2011). The *ζ*-potential of EGFR is ≈ -9.24 mV Chen et al. (2019). The electrophoretic forces alone would make the EGFRs move with a velocity of -1.5 nm/s (toward the anode, Eq. S1). However, the additional presence of the electroosmotic flow makes the negatively charged EGFRs accumulate cathodaly with an electromobility velocity of 6 nm/s (Fig. 2a, Eq. 3).

**Figure 2:**
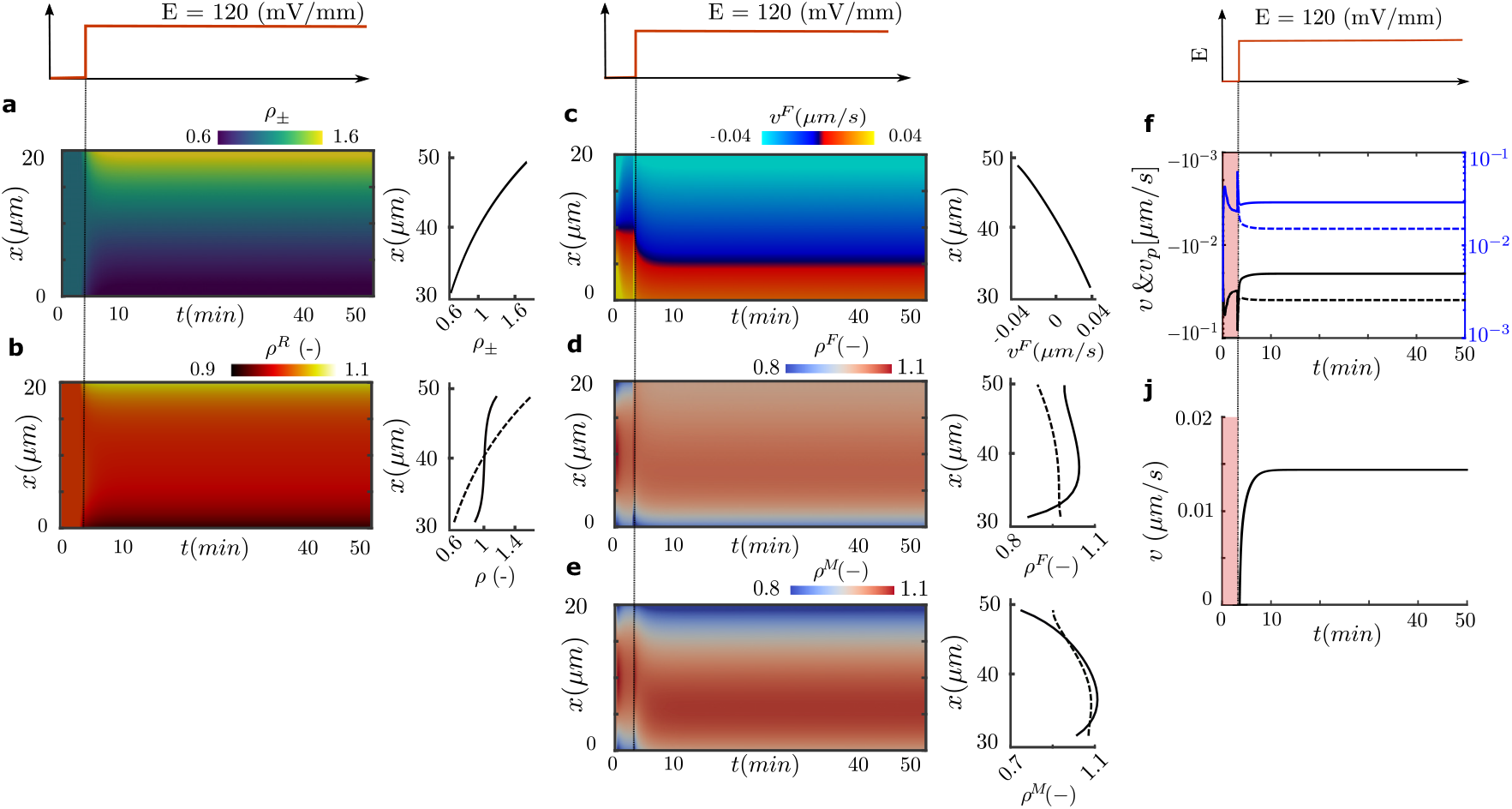
Computational results of a cell stimulated with an EF of 120mV/mm with *ζ*-potential difference of 50.76 *mV* : Kymograph of the CMPs polarization (a) and intracellular signals (b). At a steady state (right), the CMPs and the intracellular signals (responder, solid line, and activator, dash line) are shown; Kymographs of the retrograde flow (c), actin (d), and myosin (e) densities. At steady state, the retrograde flow and actin (F-actin, solid line, G-actin, dash line) and myosin densities (bound, solid line, unbound, dash line) are shown on the right; Retrograde flow (solid), blue and black at the cathodal and anodal sides of the cell, respectively, and polymerization velocity (dash), black and blue at the cathodal and anodal sides of the cell, respectively (f); Migration velocity of the cell (j). Color bars are differentiated by transmembrane, internal signal and actomyosin network variables.

The polarization of EGFRs induces a cascade of intracellular signals that we account for with a LEGI model. First, the accumulation of EGFR at the cathodal side activates PI3K pathways, which activates the responders (e.g., small GTPases Rac1 and Cdc42) (Fig. 2b). This signaling process has two main effects on the migration behavior of the cell. First, the actin polymerization velocity increases in those regions where EGFR, and consequently PI3K, Rac1, and Cdc42, accumulate. The polarization of Rac1 induces an equal polarization but in opposite directions in RhoA Nimnual et al. (2003); Ohta et al. (2006) and, consequently, in myosin activity, which results in a backward retrograde flow (Fig. 2c). The retrograde flow drags the actomyosin network, which further increases the front-rear polarization in a positive feedback loop (Fig. 2d-e). The balance between the actin polymerization velocity and the retrograde flow velocities at the cell fronts establishes the directed migration of the cell to the cathode (Fig. 2f) with a steady-state velocity of 0.014 *µ*m/s, in agreement with the velocity and direction of cell migration Zhao et al. (1999, 2002a) (Fig. 2j).

In summary, our model integrates all modules involved in the cascade of events that triggers electrotaxis. It pinpoints the fundamental role of the *ζ*-potential in the response of CMPs to an EF (EGFR in this case), describes the role of the polarization of the CMPs in the downstream regulation of PI3K and GTPases and the consequent polarization of the intracellular forces that control cell migration. The model reproduces the polarization of EGFR, PI3K, and small GTPases toward the cathode as previously described Zhao et al. (1999, 2002a); Wang et al. (2011).

### *ζ*-potential differences explain opposing electrotactic directions

As we showed above, the polarization of EGFR and the downstream signaling events polarize the cell toward the cathode. Although electrotaxis is ubiquitous across cell types and certain cell types migrate toward the cathode, others do it toward the anode as we described in the introduction. Based on an EGFR-centered model, as the main CMP involved in the sensing process, this would not be possible. However, there are other CMPs involved in downstream signals of the migratory machinery of the cell. Different CMPs, with different physical properties, have different *ζ*-potentials and sizes. Indeed, the sign of the *ζ*-potential difference should explain the polarization of the CMPs toward the cathode or the anode (see Eq. 3-4) and, consequently, the cathodal and anodal migration, respectively.

Therefore, we analyze the effect of the *ζ*-potential differences in electrotaxis by considering hypothetical cells with a *ζ*-potential difference in a range of -60 to 60 mV exposed to a range of EFs between 50 and 500mV and compute the electromotility velocity of the CMPs (Fig. 3a). These CMPs can then be responsible for the activation of downstream signals, as described for EGFR above. Electromotility velocities increase as we increase the strength of the EF and the magnitude of the *ζ*-potential difference. Increasing values of *ζ*-potential differences and EF induce an increasing accumulation of CMPs toward the cathode (*ζ*_1_ − *ζ*_2_ > 0) or anode (*ζ*_1_ − *ζ*_2_ < 0).

**Figure 3:**
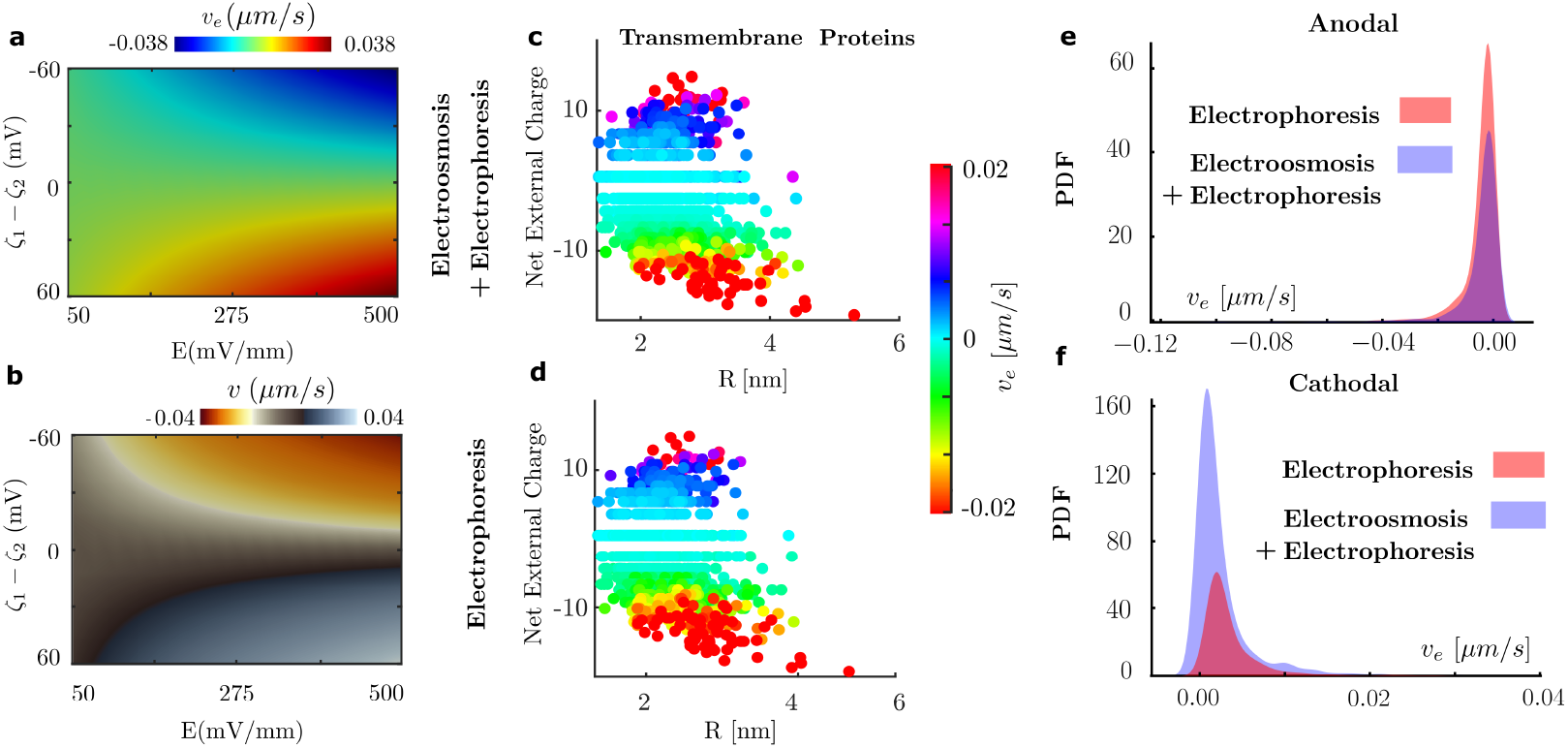
Color map of the model results for (a) electromotility and (b) migration velocities. The results are computed for a *ζ*-potential difference in a range of -60 to 60 *mV* and an EF stimulus in a range of 50 to 500 *mV*. (c) Results of electromigration velocities on transmembrane proteins when electroosmosis and electrophoresis are active and (d) when electroosmotic flows have been canceled. (e-f)For those proteins that electromigrate toward the cathode and the anode, we compute the PDF for the control (electro-osmosis + electrophoresis) and the modified system (only electrophoresis)

Then, we compute the cell migration velocity for each combination of *ζ*-potential and EFs at steady-state. As expected, our results show a clear correlation between the sign of the *ζ*-potential difference and the direction of cell migration and between the magnitude of the *ζ*-potential difference and of the EF and the migration velocity of the cell (Fig. 3b), as repetitively shown in the literature. Therefore, cells crowded by CMPs that result in *ζ*-potential differences of the same magnitude but opposite signs induce migration velocities of the same magnitude but opposite direction, i.e cathodal and anodal electrotaxis (Fig. 3b).

As one could have predicted, the model results suggest that the *ζ*-potential difference of cells, and consequently the electromotility of CMPs, may be responsible for the different electrotactic responses across cell types.

### Track down CMPs responsible for electrotaxis

Then, we wonder what specific CMPs could match a specific *ζ*-potential difference and, consequently, could be responsible for anodal or cathodal migrations. To tackle this idea, we used a previous screening of the zebrafish keratocytes proteome to describe the electromigration of CMPs (see Sarkar et al. (2019) for a complete description of the screening process and the physical properties of the proteins).

We plot the electromotility velocity *υ*_*e*_ as a function of the charge and equivalent radius of transmembrane proteins (Fig. 3c) and integral and GPI-anchored proteins (Fig. S4c-d). Our results show that 47.2% of transmembrane proteins, 46.6% of extracellular integral membrane proteins, and 45.6% of the GPI-anchored proteins redistribute toward the anode (Fig. S6). All proteins that accumulate toward the anode have net negative extracellular charges, with averaged mean charges of -7.59, -13 and -12, for transmembrane, integral, and GPI-anchored proteins, respectively. For these proteins, electrophoretic forces are larger than the drag forces due to the cathodal electroosmotic flow. For cathodal accumulating proteins, 50.5% of transmembrane proteins, 71.4% of extracellular integral membrane proteins, and 50% of the GPI-anchored proteins have positive charges, meaning that electrophoretic and electroosmotic forces cooperate to drag proteins toward the cathode. In the cases with negative CMPs, the electroosmotic flow dominates, and the CMPs polarize toward the cathode. All proteins analyzed are provided as *Dataset S1* with their physical properties, electromigration velocities. A statistical analysis is also shown in Fig. S6.

Then, we identify proteins that may be implicated in downstream signals for cell migration. For those proteins that electromigrate toward the cathode, we found proteins participating in the EGFR transactivation/metalloproteases pathway (alpha-2A adrenergic receptor) and PI3K signaling (tyrosine-protein kinase STYK1), epithelial-mesenchymal transition (lysophosphatidic acid receptor 5b), integrin involvement (opsin-5), and G protein-coupled receptors/Rho GTPases (sphingosine 1-phosphate receptor 1). In the list of anodal accumulating trans-membrane proteins, we also found proteins participating in interrelated pathways that include EGFR transactivation/metalloproteases (type-1 angiotensin II receptor) and PI3K signaling (adiponectin receptor protein 2), epithelial-mesenchymal transition (TM2 domain-containing protein 1 precursor), integrin involvement (nuclear envelope integral membrane protein 2), G protein-coupled receptors/Rho GTPases (beta-2 adrenergic receptor), and metalloproteases (tetraspanin-12). Each CMP and the affecting signaling pathway is available in *Dataset S2*.

However, the assumption that all CMPs may be dragged by the maximum electroosmotic flow velocity could be overestimated. Some proteins could have the core domains below that maximum velocity while others could have them above the debye length. Moreover, solutions of higher viscosity than the value used here could create a much slower electroosmotic flow. To analyze these scenarios, we recompute the electromigration of all CMPs without the electroosmotic flow (Fig. 3d, Fig. S4e-f). Our results show a preferential polarization of CMPs toward the anode because the electroosmotic flow, assisting the cathodal accumulation, is now canceled. Now, the percentage of CMPs accumulating toward the anode for transmembrane proteins, integral proteins, and GPI anchored proteins is 72.9%, 61%, and 72%, respectively (Fig. S6). We also plot the Probability Density Function (PDF) of proteins accumulating in the cathode and the anode for these cases (Fig. 3e-f, Fig. S4e-h). Finally, we look into CMPs that electromigrated to the cathode in the control case but switched to the anode when the electroosmotic flow is removed. We find 7 of such proteins, such as the G-protein coupled receptor 26 and 78 and urotensin-2 receptor-like, that are implicated in different pathways of cell migration (highlighted in yellow in *Dataset S2*).

Together, these results show simultaneous anodal and cathodal accumulation of transmembrane proteins involved in downstream signaling pathways of the motile machinery of the cell. Whether anodal or cathodal electrotaxis prevails will depend on the polarization of CMP, the following competition of signaling pathways, and the strength of these mechanisms that make either the anodal or cathodal electrotaxis win.

### Intracellular signaling may explain opposing migration directions

Then, we wonder how this opposing redistribution of CMPs and, consequently, multiple stimuli for intracellular signaling impacts electrotaxis. To analyze this scenario, we increase the complexity of the signaling model. We adopt previous models (see Dawes and Edelstein-Keshet (2007); Marée et al. (2012); Holmes and Edelstein-Keshet (2016) for details), which include an explicit description of small GTPases of the rho family (Rac1, Cdc42, and RhoA), phosphoinositide kinases (PI3K and PI5K), and phosphoinositides (PI(4)P, PI(4,5)P_2_, and PI(3,4,5)P_3_, or PIP, PIP2, and PIP3, respectively). This approach allows us to impose stimuli for each pathway (rho GTPases, phosphoinositide kinases, and phosphoinositides) and analyze the feedback loops between them (Fig. 4a). A complete revision of these interactions and the full coupled system of Partial Derivative Equations (PDEs) is described in *SI Appendix*.

**Figure 4:**
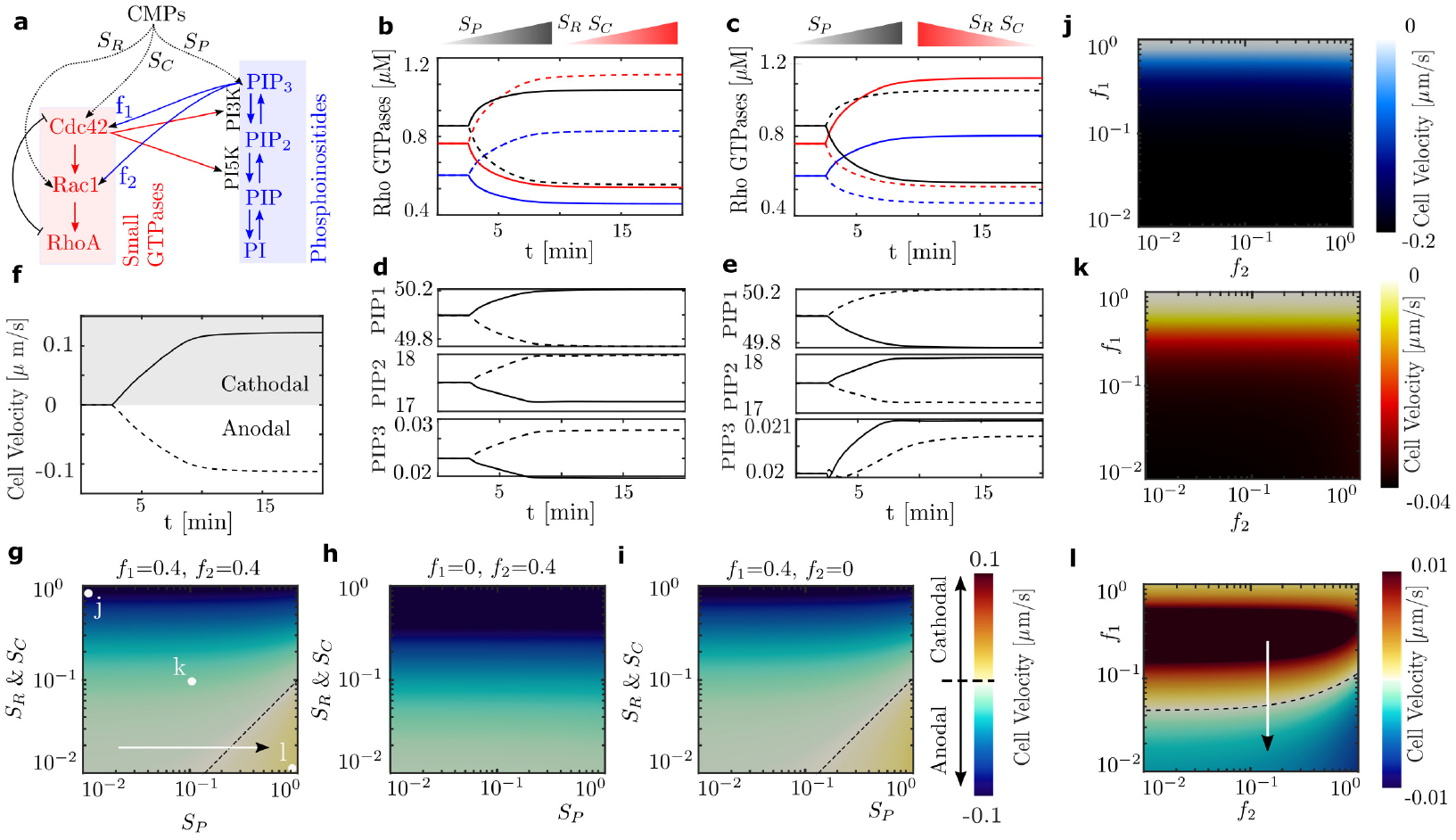
(a) Schematic representation of the signaling interactions. Arrows indicate positive feedback (activation) from one component to another. Tails indicate negative feedback (inactivation) of one component by another. *f*_1_ and *f*_2_ are the magnitudes of feedback for activating Cdc42 and Rac1 by PIP3. *S*_*i*_ is the strength of the signal from receptors to GTPases or PI3K. (b-f) Two cases are shown, where the gradient of receptors that activates PI3K is always polarized toward the cathode (grey gradient) and the polarization of receptors that activate Rac1 and Cdc42 are toward the cathode (b-c) or the anode (e-f). Evolution in time of the GTPases (b-c) and PIPs (e-f) in the cathodal (dash) and anodal (solid) side of the cell are presented. (d) Cell migration velocity for the cases presented in (b-c) and (e-f). For the case presented in (b-c), we vary the strength of the stimulus *S*_*P*_, *S*_*R*_, and *S*_*C*_ in a normal feedback loop from PIP3 to Cdc42 and Rac1 (g), when no feedback loop to Cdc42 (h) and no feedback loop to Rac1 (i). For the three cases presented in (g) in points (j-l) we vary the strength of the feedback loop from PIP3 to Cdc42 and Rac1.

In short, the model considers the GTPase’s interaction that switches between membrane-boundand cytosolic states, assuming that cycling between membrane and cytosol is very fast Jilkine et al. (2007). The GTPases are assumed to be in two forms active (*G*) and composite inactive (*G*^*i*^) given by

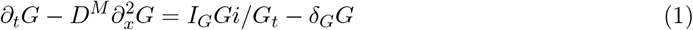

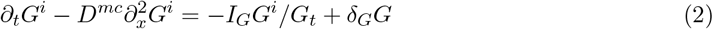

where *G* can be Rac1, Cdc42, or RhoA, and *G*_*t*_ represent the total concentration of each substance. *D*^*mc*^ is the diffusion constant that averages the respective rates of diffusion of inactive GTPase forms by the time spent on the membrane versus the cytosol and *D*^*M*^ is the diffusion constant of the active form. *δ*_*G*_ is the GAP-mediated inactivation rate and *I*_*G*_ is the GEF-mediated activation rate. The interaction between phosphoinositides, PIP, PIP2, and PIP3, are also modeled by a set of similar PDEs with reaction kinetics. Interactions between GTPases and PIPs are included in coupling terms between these sets of PDEs through kinetic reactions.

Because we don’t know all CMPs involved in the signaling process and how strongly they respond to the polarized CMPs, we explore mathematically different possibilities. We compute the polarization of two CMPs with a *ζ*-potential difference of 10 and -10 *mV* under the stimulus of an EF of 120*mV/mm* (Fig. 4), which produces anodal and/or cathodal accumulation of CMPs. Some of these CMPs can induce the activation of PI3Ks and others can activate Rac1 and Cdc42 through GEFs (Fig. 4a). Then, the initially constant activation rates *I*_*R*_, *I*_*C*_, and *k*_*pi*3*k*_ are multiplied by the resulting normalized spatial density of the three different types of CMPs and by different strength factors, *S*_*R*_, *S*_*C*_, and *S*_*P*_, to describe the effective downstream activation (Fig. 4a). Therefore, the stimulus becomes *I*_*G*_ ← *S*_*G*_*ρ*_*±*_*I*_*G*_.

We investigate possible cooperative and competing combinations of these stimuli. We take the CMP that polarizes toward the cathode to activate PI3K which, unless stated otherwise, is maintained in all our simulations. In our control case, we assume that another CMP, which also polarizes toward the cathode, activates Rac1 and Cdc42 (Fig. 4b-c). The accumulation of Cdc42 in the cathodal side of the cell induces a strong actin polymerization, that pushes the membrane forward, and an accumulation of RhoA at the anodal side, that activates myosin activity and a cathodal-to-anodal retrograde flow (Fig. S1). Eventually, the chain of cooperative positive feedback loops induces an electrotactic response toward the cathode with a migration velocity of 0.12 *µ*m/s

(Fig. 4f). Next, we consider that the CMPs responsible for activating Rac1 and Cdc42 polarize toward the anode, competing with the downstream signals of the cathodal PI3K polarization (Fig. 4d-f). Our simulation shows that cells would migrate anodally with a velocity of -0.115 *µ*m/s (Fig. 4d). These results indicate that direct activation of GTPases (Cdc42 and Rac1) and not PI3K seems to dominate the final polarization of the cell (Fig. 4d-f, Fig. S2-S3).

Then, to understand the behavior of different cell types, we wonder how different values of the strength of the stimuli (*S*_*R*_, *S*_*C*_, and *S*_*P*_) and the feedback loops from PIP3 to the GTPases (f_1_ and f_2_, Fig. 4a) impact electrotaxis (Fig. 4g-i). We focus on the case where the CMPs responsible for PI3K and Cdc42 polarize in opposite directions (Fig. 4e-f). First, we vary the stimuli strength and keep the feedback from PIP3 toward Rac1 and Cdc42 (Fig. 4g). Our results show that migration occurs preferentially toward the anode or, in other words, toward the Cdc42 polarization. However, cells migrate cathodically for ratios of *S*_*P*_ */S*_*C*_>10 following the PI3K polarization. If the feedback from PIP3 to Cdc42 is canceled, cells migrate anodally with higher velocity because the opposing stimulus coming from the PI3K polarization is inhibited (Fig. 4h). Further, the inhibition of the feedback from PIP3 to Rac1 does not affect the migration velocity (Fig. 4g,i), indicating the effect of Rac1 from the PIP3 activation is weak. These results suggest that changes in the activation strength of PIP3 have little effect on cell migration velocity and that activation of Cdc42 dominates over the activation from PI3K and Rac1 (Fig. 4g-i).

To further understand how the feedback from PIP3 to Cdc42 and Rac1 may affect the signaling process Dawes and Edelstein-Keshet (2007); Marée et al. (2012); Holmes and Edelstein-Keshet (2016), we analyze the effect of their strength (Fig. 4j-l). We analyze 3 cases (see dots in Fig. 4g) with different ratios of the stimuli strengths, *S*_*R*_, *S*_*C*_, and *S*_*P*_. As shown before, the feedback strength from PIP3 to Rac1 does not affect the migration velocity of the cell for ratios *S*_*R*_*/S*_*P*_ that initially induced anodal migration (Fig. 4j-k). If the feedback strength from PIP3 to Cdc42 is enhanced, the migration velocity also increases but doesn’t change the migration direction (Fig. 4j-k). However, for ratios of *S*_*R*_*/S*_*P*_ that induce migration toward the PI3K accumulation (point l, Fig. 4g), the migration direction strongly depends on the feedback strength from PIP3 to Cdc42 (Fig. 4l). Inhibition of the feedback from PIP3 to Cdc42 will shift the migration direction toward the anode, where Cdc42 polarizes, by recovering the effect of the direct activation of the GTPases. Together, these results unveil the complex connections between the stimuli induced by the polarized CMPs and the strength of the downstream signaling pathways. Our results show how different strengths of PIPs to GTPases feedback loops and of the stimuli induced by the CMPs electro-migration can induce anodal and cathodal electrotaxis.

### Electrotactic control by cellular manipulation

Next, we wonder how we could manipulate cells to engineer a specific response to enhance, arrest, or shift the electrotaxis direction. To do so, we look back into the main ingredients involved in electrotaxis. Cell motility can be directly modified by myosin activity, actin polymerization, or adhesion manipulations, e.g. blebistatin, CK-666, or talin knockouts, respectively. We showed through mathematical models the effect of this intracellular manipulation on cell migration (Betorz et al. (2023); Sáez and Venturini (2023)). However, these are not unique to electrotaxis.

Electrotaxis must be directly coordinated by the polarization of CMPs and the downstream signaling pathways. Therefore, one could induce changes in the electromigration direction of CMPs. The literature has proposed different ways of such manipulation, which become clear when we look at Eq. 3. One option is to modify the viscosity of the medium, which reduces the electroosmotic flow Kobylkevich et al. (2018), as demonstrated above (Fig. 3). Another option is to change the charge amount of the CMPs by modifying the pH of the medium Allen et al. (2013); Sarkar et al. (2019) or by glycosylation. Glycosylation has been shown to shift the cathodal electromigration of CMPs to the anodal Sarkar et al. (2019). For small negative values of cell surface *ζ*-potential, the electroosmotic flow due to the negative cell surface charge pushes the CMPs toward the cathode. When the CMP charge becomes more negative, presumably the negative charge generates a larger electrophoretic attraction towards the anode and, therefore, it can compensate for the drag forces toward the cathode. However, the effects on the migration of CMPs by such putative electrophoretic force are complicated by the electroosmotic effect and the *ζ*-potential variation, making its mathematical modeling of this particular aspect unfeasible.

Finally, we can also infer the signaling pathways to modify electrotaxis. A widely reported manipulation consists of blocking PI3K expression Zhao et al. (2006); Meng et al. (2011), which breaks the PIP2 to PIP3 activation and, consequently, feedback to Rac1 and Cdc42. However, there are reports of electrotactic inhibition while other works described no change in the electrotaxis response or even a shift in the migration direction Sun et al. (2018, 2023). Our model conceptualizes all these observations (point j, Fig. 4j-l). Let’s assume a cell that shows Cdc42 polarization and migration toward the cathode but PIP3 polarization is toward the anode (Fig. 4d-f). Migration inhibition would be consistent with a system in which a strong GTPases polarization does not receive enough feedback activation from PIP3, whether PI3K sufficiently activates PIP3 or not (point j, Fig. 4g,j). These results suggest the feedback from PIP3 to Cdc42 as a key mediator in inhibiting or enhancing electrotaxis. There are two options to explain reversing electrotaxis direction. We observe electrotaxis toward the Cdc42 accumulation in a system where the activation of GTPases and PI3K is weak. If we then enhance PI3K activation, which then directly feedback to PIP3, Cdc42, and Rac1, we would overrule the direct Cdc42 activation at the anodal side and promote cathodal accumulation (Fig. 4g, white arrow). Second, in such manipulated or endogenous cases of strong PI3K activation (point l, Fig. 4g), we could again reverse the migration direction either by inhibiting the PIP3 feedback to Cdc42 (Fig. 4l, white arrow). It is key to note that, when the polarization of all CMPs is aligned toward the same side of the cell, we cannot find any signaling manipulation that can reverse migration direction.

## Discussion

The control on-demand of cell migration has huge implications in cancer, wound healing, and tissue engineering. Among other external cues, an electric field represents a stable and programmable signal to control cell motility Zajdel et al. (2020). Although electrotaxis happens in a multitude of cellular systems, it is expressed with important differences. Some cell types migrate toward the anode while others do it toward the cathode. To precisely control electrotaxis, we must understand the fundamental mechanisms that enable electrotaxis across cell types.

By adopting a minimal model of cell migration Betorz et al. (2023), and extending it with electromigration of cell membrane receptors and signaling pathways implicated in cell motility, we have proposed a complete computational model of electrotaxis. First, we wondered about what membrane proteins may be involved in electrotaxis because, as previously suggested, the polarization of CMPs plays a key role in cell migration Allen et al. (2013); Sarkar et al. (2019). To answer this question we looked into the zebrafish’s proteome and identified transmembrane proteins involved in the downstream signaling pathways of electrotaxis. Our results indicate that individual CMPs electromigrate cathodally and anodally and that they may predominately accumulate anodally if electroosmotic flows are reduced, or if the charge amount is increased. For example, EGFRs and G-protein coupled receptors, which are implicated in cell motility, show cathodal polarization in a control set-up but can migrate anodally if the electroosmotic flow is canceled.

But what could happen if, as we demonstrate, multiple stimuli compete to determine the final cell polarization? With our computational model, we showed that the opposed polarization of membrane receptors can trigger downstream competing signaling pathways and that, depending on where and at what strength receptors polarize, the prevailing signal will make cells migrate anodally or cathodally. This activation strength can be cell-type dependent, and the downstream signaling competition between different activation pathways may explain antagonistic responses in electrotaxis. Our results, in addition to replicating previous experimental findings, provide a mechanistic rationale for how different cell types may exhibit anodal or cathodal migration.

These conclusions lead us to propose future experimental work, that lies beyond the scope of this work. Most of the data today usually reduces to one specific receptor and, in some cases, the studies extend to the analysis of one other intracellular signal, for example, PIP3 or PI3K. First, a detailed proteome analysis of each cell line should be performed, followed by the electromotility study of each protein and their corresponding downstream signaling pathways for cell migration. Based on our analysis, differences in cellular responses, such as migration velocities or time to respond to an electric stimulus, should respond to differences in activation strengths and the feedback loops between GTPases and PIPs. We strongly suggest that the next steps should focus on a consistent analysis of how all these signals (Rac1, Cdc42, RhoA, PI3K, PI5K, PIP, PIP2, and PIP3), and not only one of them, polarize under the EF. This extensive data analysis would tell us how these feedback loops occur, which should uniquely determine the electrotactic mechanics for a specific cell line, providing a case-by-case answer to the response of cells to an external EF.

Our computational model stands as a potential tool to guide future experimental research in electrotaxis. To this end, we present an online platform for the scientific community (uploaded in the MATLAB Central File Exchange) that, upon providing the physical properties of the membrane proteins and the electric field strength yields the distribution of the proteins, the Rho-GTPases, PIPs, and phosphoinositides, the myosin and actin distribution and the migration velocity of the cell.

Our model reduces and rationalizes electrotaxis in terms of the final polarization of the actomyosin network. However, other mechanisms could also be involved in electrotaxis. An EF also activates voltage-gated ion channels. The cell membrane depolarization would lead to the elevation in intracellular ion concentration. Ca^2+^, Na^+^, and K^+^ channels have been directly implicated in electrotaxis Nakajima et al. (2015); Djamgoz et al. (2001). Indeed, one of the immediate effects in terms of cellular response to EF stimulation is the increase in intracellular Ca^2+^ Cho et al. (1999); Mycielska and Djamgoz (2004); Zhang et al. (2018). Even though some previous data have ruled out this contribution to electrotaxis, it would be interesting to develop a computational model to analyze these cases and propose possible sensing and transductive mechanisms of electrotaxis through such electrochemical regulation events. We have also assumed that CMPs electromigrate in isolation. However, multiple CMPs, including EGFR, may coalesce to form rafts with a total negative charge larger than the *ζ*-potential of the cell membrane which would result in a negative *ζ*-potential difference. In that case, the stronger electrophoretic attraction will result in anodal migration, which has been reported in literature Lin et al. (2017). Perhaps, there are even other mechanisms that we have not yet thought about and, therefore, the experimental and theoretical work on electrotaxis will still need to work together.

## Model

To model the cascade of events toward electrotaxis, we consider a one-dimensional computational model. The cell domain, Ω, is a continuum segment with moving coordinates *x*(*t*) ∈ [*l*_*a*_(*t*), *l*_*c*_(*t*)], where *l*_*a*_(*t*) and *l*_*c*_(*t*) represents the anodal and cathodal facing boundaries of the cell and, therefore, the length of the cell is determined as *L*(*t*) = |*l*_*a*_(*t*) − *l*_*c*_(*t*)|. The boundary velocities will be given by *l*_*a*_(*t*) and *l*_*c*_(*t*), which are described next. The moving of both cell ends will depend on the velocity of the contractile actomyosin network and the velocity of actin polymerization at the cathodal and anodal fronts, which will eventually dictate the migration direction and velocity of the cell.

To model the cascade of events toward electrotaxis, we look first into the first cell sensors. If an EF can only be sensed outside the cell, then CMPs should be directly, uniquely, and firstly implicated in these opposing electrotactic responses. Therefore, we model the electro-motility of CMPs McLaughlin and Poo (1981); Allen et al. (2013); Sarkar et al. (2019), which are then responsible for the downstream polarization of intracellular signals. To reproduce the signaling layer of the process we based on previous models from literature Dawes and Edelstein-Keshet (2007); Marée et al. (2012); Holmes and Edelstein-Keshet (2016); Levchenko and Iglesias (2002); Devreotes et al. (2017); Jilkine and Edelstein-Keshet (2011). Finally, we use our previous active gel models of cell migration to computationally analyze the coupling of GTPases with migration forces of the cell under the effect of the EF Betorz et al. (2023).

### Electromotility of charged membrane components

CMPs experience two opposing forces. The drag forces from the EOF and electrophoretic attraction redistribute CMPs asymmetrically along the cell membrane McLaughlin and Poo (1981); Allen et al. (2013); Sarkar et al. (2019), generating a cathode-anode axis of polarity in cells. To measure the rate and intensity of this redistribution due to the electrophoretic and electro-osmotic mobilities of the CMPs, we use the ‘tethered sphere model’ to simplify the geometry of CMP McLaughlin and Poo (1981). Balancing the forces acting on the CMPs (*SI Appendix*), the total electro-motility velocity (*υ*_*e*_) is given by the expression:

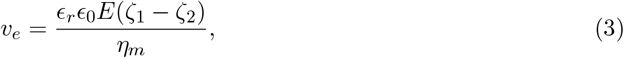

where *η*_*m*_ represents the viscosity of the cell membrane.

To describe the distribution of positive and negative CMPs, *ρ*_*±*_, in space and time, we use a convection-diffusion equation:

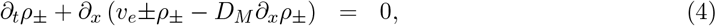

where *D*_*M*_ is the diffusion coefficient for CMPs. We assume zero Neumann boundary conditions at both ends for *ρ*_*±*_ to consider that no CMP can enter or leave the cell membrane. We also consider a normalized homogeneous initial condition *ρ*_*±*_(*x*, 0) = 1.

### Polarization of signaling cues

To model cell signaling, we first follow a Local Excitation Global Inhibition (LEGI) model Levchenko and Iglesias (2002), which represents one of the simplest yet most comprehensive mathematical models for cell signaling Devreotes et al. (2017); Jilkine and Edelstein-Keshet (2011). The LEGI model proposes PI3K and PTEN as activator (A) and inhibitor (I), respectively, of a response element (R), which represents small GTPases. Specifically, we assume R to be associated with Rac and Cdc42, which activates the cell front through actin filament polymerization. Because Rac1 and Cdc42 are mutually exclusive to RhoA Nimnual et al. (2003); Ohta et al. (2006), we assume the density of Rho A, 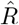, to be equal but in the opposite direction to *R*. The LEGI model accounts for all the main signaling aspects summarized in the introduction while simplifying many of the interactions and it is described by the following system of PDEs Kutscher et al. (2004):

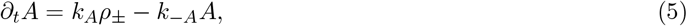

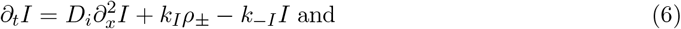

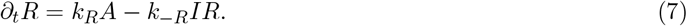

The on/off rates for each variable are denoted by *k*_*i*_ and *k*_−*i*_, respectively. The diffusion parameter for the inhibition process is denoted by *D*_*i*_. Following the hypothesis that redistribution of CMPs is responsible for electrotaxis, we take the CMP density, *ρ*_*±*_, described previously in Eq. 4, as the stimuli for cell polarization. The fast-acting local activator and the slow global inhibitor are activated in direct proportion to the temporal stimuli.

### Mechanical model of the actomyosin network

We consider an active gel model to reproduce the mechanics of actomyosin network Prost et al. (2015); Betorz et al. (2023) as

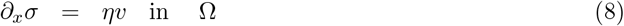

where we neglect inertial terms and assume a constitutive relation for the internal stresses,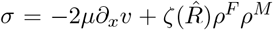, that accounts for the viscosity of the actin cortex and the actomyosin contractibility, respectively. *ρ*^*F*^ and *ρ*^*M*^ are associated with the F-actin and bound myosin motors, as described in the following sections. *υ* is the velocity of the actin network or retrograde flow in the lab frame, *µ* is its shear viscosity and *ζ* is the active contraction exerted by the contractile myosin motors that we multiplied by 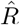 to increase contractibility in the region with a high concentration of RhoA. The term on the r.h.s. represent the friction between the moving cell and the external space, which we assume to be proportional to the cortex velocity with constant friction *η* Betorz et al. (2023); Sáez and Venturini (2023). We impose zero Neumann boundary conditions on both ends of the cell.

### Kinetics of the cell fronts

The polymerization velocity is regulated biochemically by Rac1 and Cdc42, which regulate Arp2/3 and filament assembly Pollard and Cooper (2009), and physically by the membrane tension, *τ* (*L*(*t*)), that opposes it. We assume the cell membrane tension to be determined by a linear relation *τ* = −*k*(*L*(*t*) − *L*_0_), where k is the stiffness of the cell membrane and *L*_0_ is the initial cell length.

Here, we adopt a well established model Mogilner and Oster (2003) where the tension-free polymerization velocity 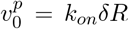 is reduced as a function of the membrane tension as 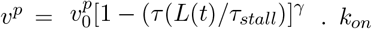 is the rate of actin assembly at the front, *τ*_*stall*_ is the tension required to stall the actin network, *γ* is a model parameter that controls the velocity decay and *δ* is the size of one single monomer at the tip of the filament. The function of 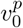 on *R* makes actin protrusions enhanced at the cell front with high levels of Rac1 and Cdc42 and inhibited at the rear.

If we know the retrograde flow velocity, *υ*, and the polymerization velocity, *υ*^*p*^, we can compute the velocity of the cell front and rear as the result of the inward retrograde flow and the outward polymerization velocity of actin against the plasma membrane. Therefore, the velocity of both ends of the cell is 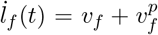 and 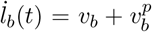, respectively. Finally, we compute the migration velocity as *V*_*cell*_ = [*İ*_*f*_ (*t*) + *İ*_*b*_(*t*)]*/*2.

### Transport of the actomyosin network

To model the actomyosin network density, we assume for simplicity a filament (F-actin), *ρ*^*F*^ (*x, t*), and a monomeric (G-actin), *ρ*^*G*^(*x, t*), form (see details in Betorz et al. (2023) and references therein). We use a coupled convection-diffusion model to describe the transport and turnover of actin in the cell:

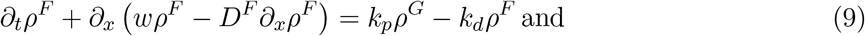

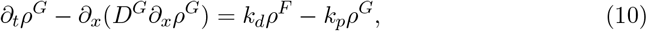

where *D*^*F*^ and *D*^*G*^ are the diffusion parameters for the F-actin and the G-actin, respectively. *k*_*d*_ and *k*_*p*_ are the depolymerization and polymerization rates. We assume that the diffusivity of the G-actin dominates and the convective term can be neglected. We define the retrograde flow in the cell frame as *w* = *υ* − *V*_*cell*_. We impose zero flux boundary conditions at both cell ends to describe that no actin form can enter or leave the cell membrane.

We also consider a two-species model of myosin motors, a form bound to the F-actin network, *ρ*^*M*^, and an unbound form, *ρ*^*m*^. The redistribution of *ρ*^*M*^ and *ρ*^*m*^ is again modelled through a convection-diffusion equation as

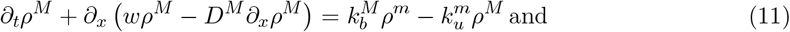

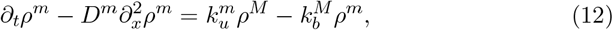

where *D*^*M*^ and *D*^*m*^ are the diffusion parameters for the bound and unbound forms of myosin, respectively.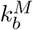 and 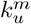 are the binding and unbinding rates of myosin to the F-actin. Again, we assume that the diffusivity of the unbound myosin dominates the convective term. We also assume zero-flux boundary conditions at both cell ends, to describe that no myosin form can enter or leave the cell membrane.

### Numerical solution of the problem and model parameters

We solve computationally the coupled system of Eqs. 4-8 and 9-12 in a staggered way. We use the Finite Element Method (FEM) to discretize the system of equations in space and an implicit second-order Crank-Nicholson method to discretize the parabolic equations in time Donea and Huerta (2003). The non-linear system of equations (Eqs. 7) is solved iteratively using the Newton-Raphson method.

The FE solution of the parabolic equations will oscillate if *Pe* > 1 (i.e. if the problem becomes convective dominant). As the convective velocity, *w*, is not known a priori, we cannot guarantee that *Pe* < 1. Therefore, we include a Stream-Upwind Petrov Galerkin (SUPG) stabilization term to keep the number of elements of our domain constant and overcome numerical oscillations in our solution Donea and Huerta (2003); Betorz et al. (2023).

Unless specified otherwise, all model parameters used in the simulations are summarized in Table S1.

### Proteomic analysis

To analyze the proteins involved in electrotaxis, we took a previous proteomic analysis of zebrafish keratocytes (see Sarkar et al. (2019) for details). In short, the zebrafish proteome was downloaded from the NCBI website. The proteins were organized into transmembrane, integral, and GPI-anchored proteins, and physical properties were added to the data set. Then, we complete the data set with the computed electromigration velocity and direction. For transmembrane proteins, we separated the proteins accumulating toward the anode and the cathode, where we screened them through STRING Database User Survey Szklarczyk et al. (2023) to obtain those proteins more prevalent in the plasma membrane. For those selected proteins, we check their biological function in the UniProt Consortium The UniProt Consortium (2023) and the Zebrafish Information Network (ZFIN) Bradford et al. (2022) database to find those proteins directly related to signaling in cell migration, either with positive or negative effects.

## 1 Acknowledgments

PS acknowledges the support of the Spanish Ministry of Science and Innovation (Grant PID2022-142178NB-I00 funded by MCIN/ AEI/10.13039/501100011033/ FEDER, UE). S.K. was supported by the Spanish Ministry of Economy and Competitiveness (Grant number: PRE2020-095851). C.R. and F.T were supported by the Spanish Ministry of Science and Innovation (Grant PID2020-115910RB-I00). We thank Y. Takada and K. Zhu for their insightful discussions.

