## Supplementary material for "Competing signaling pathways controls electrotaxis": SI appendix

### Forces acting on the redistribution of charged membrane components

We start analyzing how CMPs electromigrate along the cell membrane. When immersed in a medium, cells are typically surrounded by electrolytes, containing mobile ions, which are either attracted or repulsed by the charged cell surface. Most cell surfaces bear a negative charge and, therefore, positive ions accumulate nearby and develop well-differentiated regions within the Electrical Double Layer (EDL). The ions surrounding the charged surface are strongly bound to it and they are considered to be immobile. This layer of ions is called the Stern layer and its thickness is of the order of the ionic radius. The electric potential on the Stern layer is referred to as the Stern potential. Ions that are located beyond the Stern plane form the diffuse mobile part of the EDL. A slipping plane, also known as a shear plane, where the no-slip fluid flow boundary condition is known to apply, separates mobile ions from those that remain firmly attached to the surface. At this plane, the electric potential is called Zeta ($\zeta$)-potential.

When an EF is imposed on a cell, the EDL is also exposed to the field. Electrostatic forces act on the ions and, when they move, they drag the fluid around them. This creates an electrically generated fluid flow known as electro-osmotic flow (EOF). The electro-osmotic mobility near a cell surface is inversely proportional to the aqueous medium viscosity and directly depends on the cell surface $\zeta$ potential ($\zeta_2$)Masliyah and Bhattacharjee (2006),

$$\mu_{eo} = \frac{v_{eo}}{E} = -\frac{\epsilon_r \epsilon_0 \zeta_2}{\eta_w} \tag{S1}$$

$v_{eo}$ is the electro-osmotic velocity, $\epsilon_r$ is the dielectric constant of the medium, $\epsilon_0$ is the permittivity of free space, $E$ is the applied Electric field strength and $\eta_w$ is the viscosity of the aqueous medium.

Embedded on the cell surface, CMPs are dragged by the EOF and, consequently can migrate along the cell membrane. Moreover, these charged particles move due to electrostatic attraction and also experience a local EOF around them. The effect of these two phenomena is referred to as electrophoresis. To quantify the velocity at which the CMPs move in the presence of an external EF due to electrophoresis, the Helmholtz-Smoluchowski equation calculates the electrophoretic mobility, $\mu_{ep}$ Lin et al. (2017); McLaughlin and Poo (1981), as

$$\mu_{ep} = \frac{v_{ep}}{E} = \frac{\epsilon_r \epsilon_0 \zeta_1}{\eta_w}, \tag{S2}$$

where, $v_{ep}$ is the electrophoretic velocity of the CMP, and $\zeta_1$ is the $\zeta$-potential of the CMP.

The electrophoretic motility can also be derived from the Einstein relation, $\mu_{ep} = zDF/RT$, where $z$ is the charge on the molecule, $D$ is the diffusion coefficient for the plasma membrane macromolecules, $F$ is the Faraday's constant, $R$ is the universal gas constant and $T$ is the temperature. In terms of the absolute charge $ze$, with $e$ the elementary charge, the Boltzmann constant $k_B$ and diffusion coefficient for a spherical particle $r$, $D = k_B T/6\pi\eta_w r$, the electrophoretic mobility is

$$\mu_{ep} = \frac{qe}{6\pi\eta_w r}. \tag{S3}$$

Comparing Eqs. S3 to S2, we can define $\zeta_1$ as

$$\zeta_1 = \frac{qe}{6\pi\epsilon_0\epsilon_r r}. \tag{S4}$$

### Signaling pathways and mathematical model

Specifically, Cdc42, Rac1, and RhoA, members of the family of Rho-GTPases and highly conserved in various eukaryotic cells, are directly involved in regulating the motility forces of the cell such as actin polymerization Pollard and Cooper (2009), myosin contractility Sun et al. (2013, 2018); Jeon et al. (2019) and cell adhesion Fukata et al. (2003). These members of the Rho family proteins alternate between an active GTP-bound state on the membrane and an inactive GDP-bound state on the membrane or in the cytosol. They activate in response to Guanine nucleotide exchange factors (GEFs) by promoting the replacement of GDP with GTP, while GTPase-activating proteins (GAPs) accelerate GTP hydrolysis, leading to protein inactivation Raftopoulou and Hall (2004); Moon (2003). When a ligand activates polarized GPCRs and RTKs, it also induces a polarized expression of GEFs, which propagates downstream to generate a polarization of Rho-GTPases. The active forms of Rho GTPases regulate the activity of numerous proteins involved in cytoskeletal processes, influencing the polymerization and depolymerization of the actin cytoskeleton Mitchison and Cramer (1996); Ridley (2001); Wedlich-Soldner et al. (2003); Chung et al. (2000). Cdc42 and Rac are highly concentrated at the leading edge Gardiner et al. (2002); Herzmark et al. (2007); Weiner et al. (1999, 2007) and Rho is predominantly localized at the rear Xu et al. (2003); Srinivasan et al. (2003); Kraynov et al. (2000); Nalbant et al. (2004); Wong et al. (2006). Rac recruits downstream effectors such as Arp2/3 and the Scar/WAVE complex. Arp2/3 is activated near the membrane by WASp or N-WASp and facilitates side-branching, nucleation of new barbed ends Svitkina and Borisy (1999); Amann and Pollard (2001); Prehoda et al. (2000); Blanchoin et al. (2000) and polymerization against the cell membrane Pollard and Cooper (2009). As a result, actin-driven membrane protrusions form at regions of high Rac activity, leading to an increase in cell surface area and inducing a global rise in membrane tension Houk et al. (2012); Goehring and Grill (2013). Similar to Rac, Cdc42 interacts with membrane-associated proteins like WASp or N-WASp, promoting the activation of Arp2/3. Cdc42 can also activate Rac Nobes and Hall (1995). Active Rac forms a gradient with the highest concentration at the leading edge of migrating Swiss 3T3 fibroblasts Kraynov et al. (2000). Similar experiments have shown high levels of active Cdc42 at cell edges undergoing remodeling Nalbant et al. (2004). Gradients of active Cdc42 and Rac, with the highest concentration at the leading edge, have been detected in motile HT1080 cells Itoh et al. (2002). Rho, on the other hand, promotes myosin light chain phosphorylation through Rho kinase (ROCK) Kimura et al. (1996); Bishop and Hall (2000); Ridley (2001). Importantly, it is believed that the distribution of active Rho is inverse to that of Cdc42 and Rac in motile cells Raftopoulou and Hall (2004); Zondag et al. (2000) thanks to mutual inhibitory feedback loops between Rac and RhoA Giniger (2002); Nimnual et al. (2003); Ohta et al. (2006). ROCK can suppress Rac activity by targeting specific Rac-specific GEFs and GAPs Tsuji et al. (2002); Worthylake and Burridge (2003); Ohta et al. (2006). Cdc42 has been also shown to have both positive and negative regulatory effects on Rho activation while Rho, in turn, acts as a local antagonist to Cdc42 Sander et al. (1998). In neutrophils, mutual exclusion between Cdc42 and Rho has been also observed Giniger (2002).

The polarized GPCRs and RTKs can also induce a polarized expression of membrane lipids, PI(4)P, PI(4,5)$P_2$, and PI(3,4,5)$P_3$ (PIP, PIP2, and PIP3), Phosphoinositide 3-kinase (PI3K) and phosphoinositide-4-phosphate 5-kinases (PI5Ks), which are involved in the directed cell migration during electrotaxis Zhao et al. (2002); Lin et al. (2017). Active Cdc42 and Rac activate PI3K Zheng et al. (1994); Srinivasan et al. (2003); Merlot and Firtel (2003) which generates PIP3 from PIP2, which becomes upregulated at the cell location closest to a stimulating signal Weiner et al. (2002); Welch et al. (2003). Rac has been also shown to interact with PI5K, which converts PIP to PIP2, to enhance their activity Tolias et al. (2000); Czech (2000). In a positive close loop, Rac activation is dependent on PI3K and its inhibition blocks Rac activation Sander et al. (1998); Rickert et al.

(2000). On the other hand, PTEN, which converts PIP3 back to PIP2, shifts to the opposite end. Indeed, PIP2 is central to cell migration because it activates actin polymerization Prehoda et al. (2000); Papayannopoulos et al. (2005) and Cdc42 and recruits Rac Rajnicek et al. (2006) which, in turn, controls cell polarity Yeung et al. (2008) and activate the PI3K-Akt pathway Zhao et al. (2006). Moreover, PIP2 inhibits the association of capping protein with the barbed ends of actin filaments Ridley (2001) and it is required to activate WASp and N-WASp, which subsequently activate Arp2/3 Rohatgi et al. (1999); Higgs and Pollard (2000); Rohatgi et al. (2001). On the other hand, PIP3 is necessary for the activation of Cdc42 and Rac Wang et al. (2002); Weiner et al. (2002); Srinivasan et al. (2003); Hawkins et al. (1995). Additionally, Rho activates Rho kinase, which activates PTEN, influencing its spatial exclusion from regions with high PI3K and Cdc42 activity Li et al. (2005). When a cell is exposed to a chemoattractant gradient, PTEN is released from the membrane at the cell's front, allowing PI3K to associate with the membrane. PTEN remains bound to the sides and back of the stimulated cell. This spatial redistribution of PI3K and PTEN leads to elevated levels of PIP3 at the leading edge Funamoto et al. (2002); Iijima and Devreotes (2002); Huang et al. (2003). Inhibiting PI3K has been shown to affect cell polarity and motility in some cases but not in others, leading to uncertainties regarding the roles of phosphoinositides in chemotaxis Jiménez et al. (2000); Rickert et al. (2000); Wang et al. (2002). It has been also shown that mammalian neutrophils and Dictyostelium discoideum cells polarize with no PI3K activity Franca-Koh et al. (2007); Ferguson et al. (2007); Nishio et al. (2007). In terms of phosphoinositides, high concentrations of PIP provide more substrate for the formation of PIP3, through increased PIP2 formation. Assuming that adaptation occurs downstream of G-protein activation, it implies that the G-protein and its effectors, including PI3K, remain elevated as long as the ligand is present.

The model takes into account all these interactions, and we make use of a series of previous models (see Dawes and Edelstein-Keshet (2007); Marée et al. (2012); Holmes and Edelstein-Keshet (2016) for further details). All model parameters and their values are summarized in the table S1. In summary, the active forms of all GTPases satisfy:

$$\frac{\partial R}{\partial t} = (I_R + \alpha C)\frac{R_i}{R_{tot}} - d_R R + D_m \partial_x^2 R \tag{S5}$$

$$\frac{\partial C}{\partial t} = (\frac{I_C}{(1+(\frac{\rho}{a_1})^n})\frac{C_i}{C_{tot}} - d_C C + D_m \partial_x^2 C \tag{S6}$$

$$\frac{\partial \rho}{\partial t} = (\frac{(I_\rho + \beta R)}{(1+(\frac{C}{a_2})^n})\frac{\rho_i}{\rho_{tot}} - d_\rho \rho + D_m \partial_x^2 \rho \tag{S7}$$

The $C_{tot}$, $R_{tot}$, $\rho_{tot}$ are the total concentrations of the Cdc42, Rac and Rho. $R_i, C_i, \rho_i$ are the concentrations of the respective inactive forms. $I_c, I_R$ and $I_{rho}$ are baseline activation rates. The values $a_1$ and $a_2$ are the Rho and Cdc42 concentrations that elicit a half-maximal drop of Cdc42 and Rho activation, respectively. The value $\alpha$ determines the rate of Cdc42-enhanced activation of Rac and $\beta$ determines the rate of Rac-enhanced Rho activation. The value $P_{3b}$ is the baseline concentration of PIP3 found in a resting cell.The inactive forms diffuse much faster ($D_m \ll D_{mc}$), they satisfy the equations:

$$\frac{\partial R_i}{\partial t} = -(I_R + \alpha C)\frac{R_i}{R_{tot}} + d_R R + D_m \partial_x^2 R \tag{S8}$$

$$\frac{\partial C_i}{\partial t} = -(\frac{I_C}{(1+(\frac{\rho}{a_1})^n})\frac{C_i}{C_{tot}} + d_C C + D_m \partial_x^2 C \tag{S9}$$

$$\frac{\partial \rho_i}{\partial t} = -(\frac{(I_\rho + \beta R)}{(1 + (\frac{C}{a_2})^n})\frac{\rho_i}{\rho_{tot}} + d_\rho \rho + D_m \partial_x^2 \rho \tag{S10}$$

The interactions between phosphoinositides and phosphoinositide kinases are also modeled mathematically as

$$\partial_t P_1 - Dp\partial_x^2 P_1 = I_{P1} - \delta_{P1} P_1 + k_{21} P_2 - f_{PI5K} P_1 \tag{S11}$$
$$\partial_t P_2 - Dp\partial_x^2 P_2 = -k_{21} P_2 + f_{PI5K} P_1 - f_{PI3K} P_2 + f_{PTEN} P_3 \tag{S12}$$
$$\partial_t P_3 - Dp\partial_x^2 P_3 = f_{PI3K} P_2 - f_{PTEN} P_3 \tag{S13}$$

where it is assumed that the conversion to PIP occurs at a constant rate, $I_{P1}$ and that all PIs diffuse in the membrane at a uniform rate $D_p$. The model also describes the fact that Rac enhances the conversion of P1 to P2 (via PI5K), of P2 to P3 via PI3K, and that Rho enhances the conversion of P3 to P2 via PTEN. The feedbacks are incorporated through the functions

$$f_{PI5K} = \frac{k_{pi5k}}{2}(1 + \frac{R}{R_b}), \tag{S14}$$
$$f_{PI3K} = \frac{k_{pi3k}}{2}(1 + \frac{R}{R_b}), \text{ and} \tag{S15}$$
$$f_{PTEN} = \frac{k_{pten}}{2}(1 + \frac{\rho}{\rho_b}). \tag{S16}$$

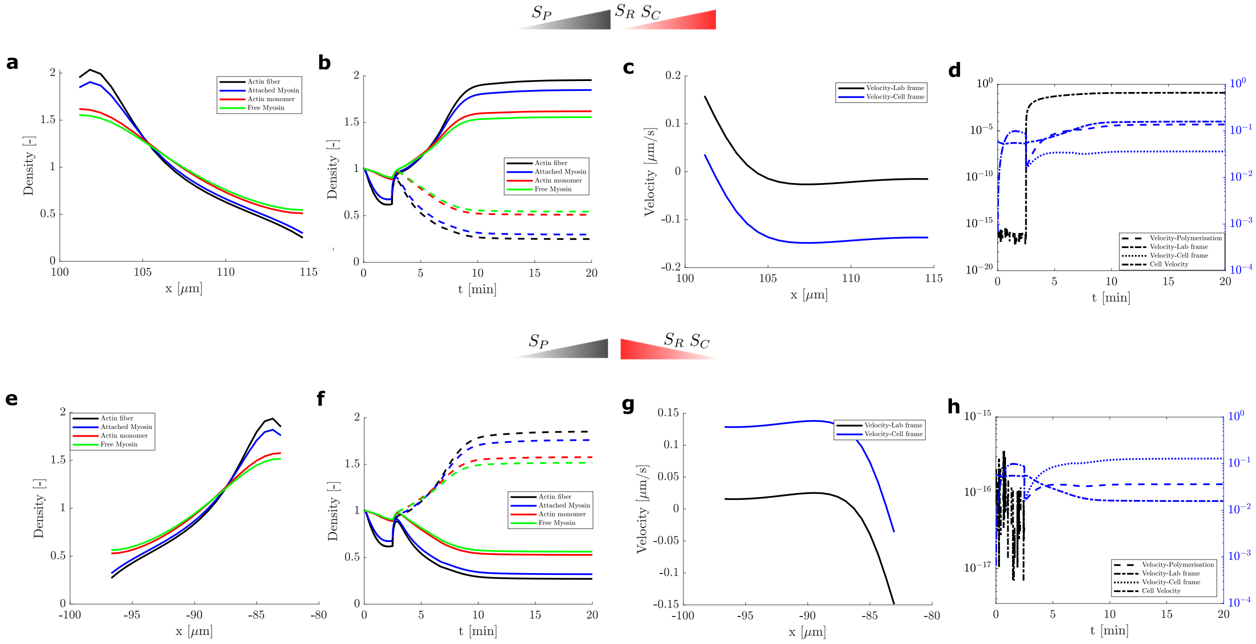


Figure S1: (a-d) Polarization of receptors that activates PI3K, Rac1 and Cdc42 are polarized toward the cathode. (e-h) The polarization of receptors that activate PI3K are polarized toward the cathode and receptors that polarize Rac1 and Cdc42 are toward the anode. (a) At steady-state, the variation of density of actin fibers (black), actin monomers (red), and bound and unbound myosin (blue and green respectively) are all concentrated towards the anode, while the time evolution is shown in (b). The spatial variation of retrograde flow velocity in the lab and cell frame are plotted in black and blue, respectively. (d) Time variation of polymerization velocity (dash), velocity of cell frame once the cell starts moving (dotted line) and cell velocity (dot-dashed line).

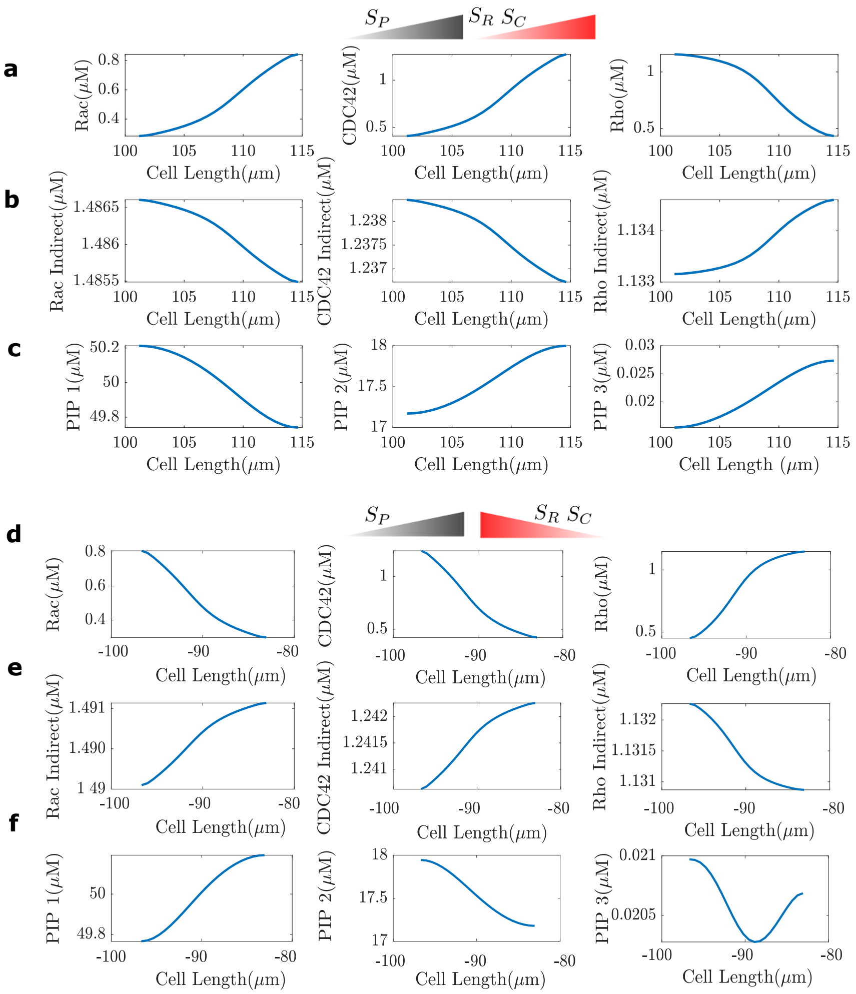


Figure S2: (a-c) Polarization of receptors that activates PI3K, Rac1 and Cdc42 are polarized toward the cathode. (d-f) The polarization of receptors that activate PI3K are toward the cathode and receptors that polarize Rac1 and Cdc42 are toward the anode. (a,d) At steady-state, the number of active GTPases, (b,e) inactive GTPases, and (c,f) PIPs is shown

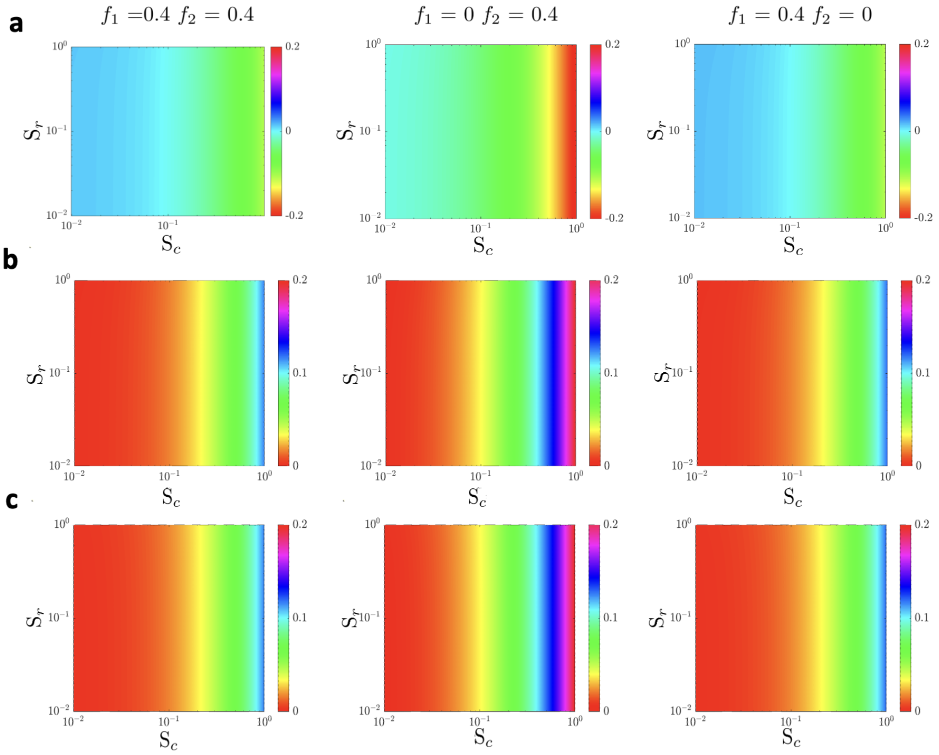


Figure S3: (a) Polarization of receptors that activates PI3K is toward the cathode, and the polarization of receptors that activates both Rac1 and CDC42 is towards the anode. (b-c) The gradient of receptors that activate PI3K is not polarized. The gradient that activates both Rac1 and CDC42 was polarized towards the cathode (b), while the gradient of Rac1 was polarized to the anode and Cdc42 was to the cathode (c). The heat-maps represent the velocity of the cell when we do a parameter sweep of the strength of stimulus of $S_r$ and $S_c$. (First column) standard case (second column) when no feedback loop from PIP3 to Cdc42 (third column) when no feedback loop from PIP3 to Rac The results show the minimal impact of the Rac1 polarization in the migration of the cell (b-c). PI3K and Cdc42 have a stronger impact on cell migration, although the effect of Cdc42 polarization is predominant.

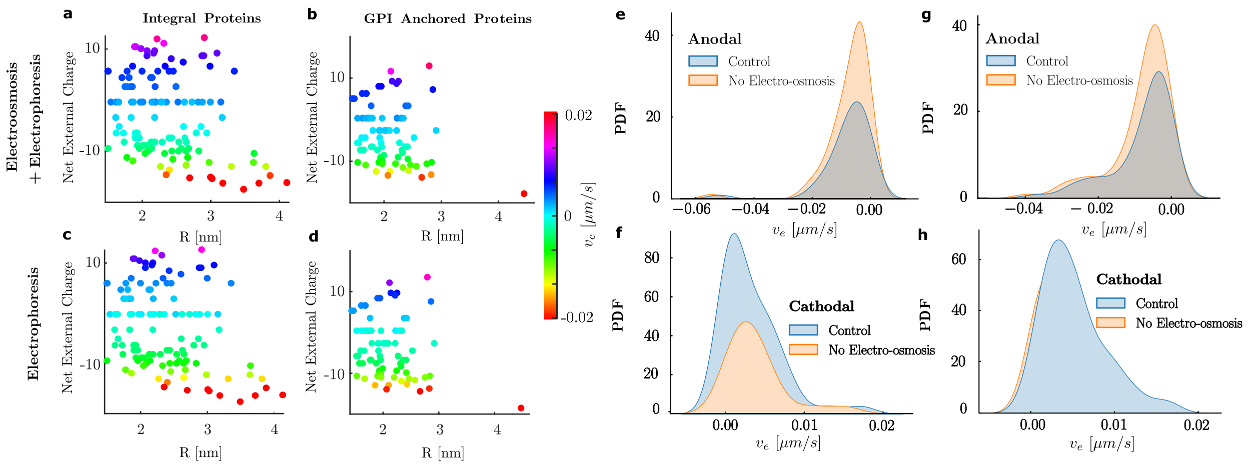


Figure S4: Results of electromigration velocities on integral (a,c) and GPI-anchored (b,d) proteins when electroosmosis and electrophoresis are active (a-b) and when electroosmotic flows have been canceled (c-d). (e-h)For those proteins that electromigrate toward the cathode and the anode, we compute the PDF for the control (electro-osmosis + electrophoresis) and the modified system (only electrophoresis) for integral (e,f) and GPI-anchored (g,h) proteins.

| Parameter | Definition | Value | Source |
|---|---|---|---|
| $k_a$ / $k_{-a}$ | The on-off rates for Activation | 0.5 | Kutscher et al. (2004) |
| $k_I$ / $k_{-I}$ | The on-off rates for Inhibition | 0.2 | Kutscher et al. (2004) |
| $k_R$ / $k_{-R}$ | The on-off rates for Rac1/CDC42 | 0.25 | Kutscher et al. (2004) |
| $L_0$ ($\mu$m) | Initial length of the cell | 10 | |
| dt | Time step for simulations | 0.4 | |
| $\eta_m$ ( $\mu$Pa.s ) | Cell Membrane viscosity | 0.2-0.3 | Houk et al. (2012); Lin et al. (2017) |
| $k_p$ / $k_d$ | Polymerization and De-polymerization rate | 0.1 | Betorz et al. (2023) |
| $k_u^m$ / $k_b^M$ | Myosin unbinding and binding rates | 0.1 | Betorz et al. (2023) |
| $D_F$ ($\mu$m/s) | F-actin Diffusion parameter | 0.3 | Betorz et al. (2023) |
| $D_G$ ($\mu$m/s) | G-actin Diffusion parameter | 3 | Betorz et al. (2023) |
| $D_M$ ($\mu$m/s) | Bound Myosin Diffusion parameter | 0.4 | Betorz et al. (2023) |
| $D_m$ ($\mu$m/s) | Unbound Myosin Diffusion parameter | 0.8 | Betorz et al. (2023) |
| $\eta$ (kPa.s/$\mu$m) | Friction Coefficient | 0.1 | Betorz et al. (2023) |
| $v_0^p$ ($\mu$m/s) | Tension free polymerization velocity | 0.55 | |
| $\mu$ (kPa.s) | Actin cortex viscosity | 10 | |
| $C_b$, $R_b$, $\rho_b$ | Typical levels of active Cdc42, Rac, Rho | 1, 3, 1.25 $\mu$M | (Marée et al. (2006)) |
| $C_{tot}$, $R_{tot}$, $P_{tot}$ | Total levels of Cdc42, Rac, Rho | 2.4, 7.5, 3.1 $\mu$M | (Marée et al. (2006)) |
| $I_C$, $I_R$, $I_\rho$ | Cdc42, Rac, Rho activation input rates | 3.4, 0.5, 3.3 $\mu$M s$^{-1}$ | (Marée et al. (2006)) |
| $a_1$ | Rho level for half-maximum inhibition of Cdc42 | 1.25 $\mu$M | (Marée et al. (2006)) |
| $a_2$ | Cdc42 level for half-maximum inhibition of Rho | 1 $\mu$M | (Marée et al. (2006)) |
| n | Hill coefficient of Cdc42-Rho mutual inhibition | 3 | (Marée et al. (2006)) |
| $\alpha$ | Cdc42-dependent Rac activation rate | 4.5 s$^{-1}$ | (Marée et al. (2006)) |
| $\beta$ | Rac-dependent Rho activation rate | 0.3 s$^{-1}$ | (Marée et al. (2006)) |
| $d_C$, $d_R$, $d_\rho$ | Decay rates of activated Rho-proteins | 1 s$^{-1}$ | (Zhang and Zheng (1998)) |
| $D_m$, $D_{mc}$ | Diffusion coefficient of active, inactive Rho-proteins | 0.1, 50 $\mu$m$^2$ s$^{-1}$ | (Postma et al. (2004)) |
| $I_{P1}$ | $PIP_1$ input rate | 10.5 mM s$^{-1}$ | (Dawes and Edelstein-Keshet (2007)) |
| $d_{P1}$ | $PIP_1$ decay rate | 0.21 s$^{-1}$ | (Dawes and Edelstein-Keshet (2007)) |
| $k_{PI5K}$ | $PIP_1$ to $PIP_2$ baseline conversion rate (by PI5K) | 0.084 s$^{-1}$ | (Dawes and Edelstein-Keshet (2007)) |
| $k_{21}$ | $PIP_2$ to $PIP_1$ conversion rate | 0.14 s$^{-1}$ | (Dawes and Edelstein-Keshet (2007)) |
| $k_{PI3K}$ | $PIP_2$ to $PIP_3$ baseline conversion rate (by PI3K) | 0.00072 s$^{-1}$ | (Dawes and Edelstein-Keshet (2007)) |
| $k_{PTEN}$ | $PIP_3$ to $PIP_2$ baseline conversion rate (by PTEN) | 0.43 s$^{-1}$ | (Dawes and Edelstein-Keshet (2007)) |
| $D_P$ | PI diffusion rate | 0.5–5 mm$^2$ s$^{-1}$ | (Postma et al. (2004)) |
| $P_{1b}$, $P_{2b}$, $P_{3b}$ | Typical levels of $PIP_1$, $PIP_2$, $PIP_3$ | 50, 30, 0.05 mM | ( Holmes and Edelstein-Keshet (2012)) |

Table S1: Parameters, values for all model equations.

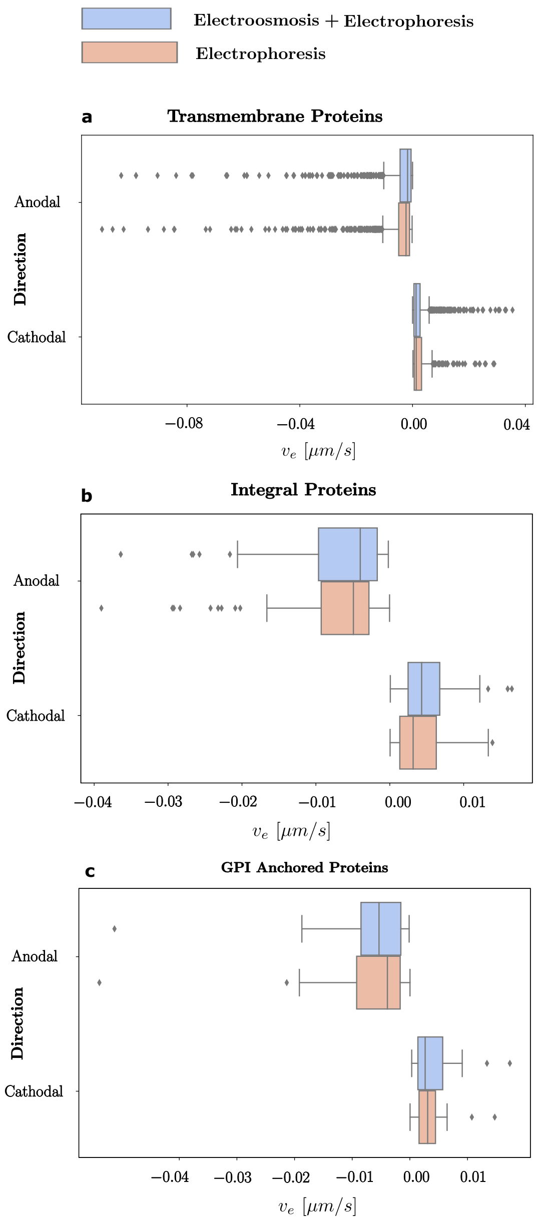


Figure S5: Boxplot of the distribution of velocities of the transmembrane (a), integral (b), and GPI-anchored (c) proteins. The control case (electrophoresis and electro-osmosis considered) is shown in blue and the case with no electro-osmosis in red. The velocity distribution shows that transmembrane proteins have a narrow range for velocities near the median. The velocity for proteins moving in the anodal direction is negative and cathodal direction is positive. Upon removal of electro-osmotic effect, the median velocity in the anodal moving proteins becomes more negative and the median velocity becomes slightly more positive for cathodal moving proteins. The trailing dots show the outlier velocities in the distribution.

| | Electrophoresis+ Electroosmosis | | | Electrophoresis | | |
|---|---|---|---|---|---|---|
| Transmembrane | | Cathodal | Anodal | | Cathodal | Anodal |
| | Mean | 0,002521 | -0,00495 | Mean | 0,002828 | -0,00548 |
| | N | 1147 | 1024 | N | 517 | 1400 |
| | Std_dev | 0,004036 | 0,073081 | Std_dev | 0,004077 | 0,009845 |
| | Median (boxplot) | 0,001242 | -0,00053 | Median (boxplot) | 0,001319 | -0,00225 |
| Integral | | Cathodal | Anodal | | Cathodal | Anodal |
| | Mean | 0,005291 | -0,00731 | Mean | 0,00403 | -0,00774 |
| | N | 63 | 55 | N | 45 | 73 |
| | Std_dev | 0,003724 | 0,008343 | Std_dev | 0,003469 | 0,0082 |
| | Median (boxplot) | 0,004313 | -0,004 | Median (boxplot) | 0,003159 | -0,00492 |
| GPI-Anchored | | Cathodal | Anodal | | Cathodal | Anodal |
| | Mean | 0,003864 | -0,00693 | Mean | 0,00377 | -0,0065 |
| | N | 44 | 37 | N | 22 | 60 |
| | Std_dev | 0,003552 | 0,00889 | Std_dev | 0,003453 | 0,008111 |
| | Median (boxplot) | 0,002652 | -0,00535 | Median (boxplot) | 0,00306 | -0,0039 |

Figure S6: Results of mean velocity, standard deviation, number of proteins, and median for the transmembrane, integral, and GPI-anchored proteins in the control case (electrophoresis and electro-osmosis considered) and the case with no electro-osmosis.

V. Papayannopoulos, C. Co, K. E. Prehoda, S. Snapper, J. Taunton, and W. A. Lim. A Polybasic Motif Allows N-WASP to Act as a Sensor of PIP2 Density. *Molecular Cell*, 17(2):181–191, Jan.

2005. ISSN 10972765. doi: 10.1016/j.molcel.2004.11.054. URL https://linkinghub.elsevier.com/retrieve/pii/S1097276504007993.

T. D. Pollard and J. A. Cooper. Actin, a central player in cell shape and movement. *Science*, 326(5957):1208–1212, 2009. ISSN 1095-9203. doi: 10.1126/science.1175862. URL http://www.pubmedcentral.nih.gov/articlerender.fcgi?artid=3677050&tool=pmcentrez&rendertype=abstract. ISBN: 0036-8075.

M. Postma, L. Bosgraaf, H. M. Loovers, and P. J. M. Van Haastert. Chemotaxis: signalling modules join hands at front and tail. *EMBO reports*, 5(1):35–40, Jan. 2004. ISSN 1469-221X. doi: 10.1038/sj.embor.7400051. URL https://www.embopress.org/doi/full/10.1038/sj.embor.7400051. Num Pages: 40 Publisher: John Wiley & Sons, Ltd.

K. E. Prehoda, J. A. Scott, R. D. Mullins, and W. A. Lim. Integration of Multiple Signals Through Cooperative Regulation of the N-WASP-Arp2/3 Complex. *Science*, 290(5492):801–806, Oct. 2000. doi: 10.1126/science.290.5492.801. URL https://www-science-org.recursos.biblioteca.upc.edu/doi/10.1126/science.290.5492.801. Publisher: American Association for the Advancement of Science.

M. Raftopoulou and A. Hall. Cell migration: Rho GTPases lead the way. *Developmental Biology*, 265(1):23–32, Jan. 2004. ISSN 00121606. doi: 10.1016/j.ydbio.2003.06.003. URL https://linkinghub.elsevier.com/retrieve/pii/S001216060300544X.

A. M. Rajnicek, L. E. Foubister, and C. D. McCaig. Growth cone steering by a physiological electric field requires dynamic microtubules, microfilaments and Rac-mediated filopodial asymmetry. *Journal of Cell Science*, 119(Pt 9):1736–1745, 2006. ISSN 0021-9533. doi: 10.1242/jcs.02897. URL http://www.ncbi.nlm.nih.gov/pubmed/16595545. ISBN: 0021-9533.

P. Rickert, O. D. Weiner, F. Wang, H. R. Bourne, and G. Servant. Leukocytes navigate by compass: roles of PI3K and its lipid products. *trends in CELL BIOLOGY*, 10, 2000.

A. J. Ridley. Rho GTPases and cell migration. *Journal of Cell Science*, 114(15):2713–2722, 2001. ISSN 00219533. doi: 10.1242/jcs.114.15.2713.

R. Rohatgi, L. Ma, H. Miki, M. Lopez, T. Kirchhausen, T. Takenawa, and M. W. Kirschner. The Interaction between N-WASP and the Arp2/3 Complex Links Cdc42-Dependent Signals to Actin Assembly. *Cell*, 97(2):221–231, Apr. 1999. ISSN 00928674. doi: 10.1016/S0092-8674(00)80732-1. URL https://linkinghub.elsevier.com/retrieve/pii/S0092867400807321.

R. Rohatgi, P. Nollau, H.-Y. H. Ho, M. W. Kirschner, and B. J. Mayer. Nck and Phosphatidylinositol 4,5-Bisphosphate Synergistically Activate Actin Polymerization through the N-WASP-Arp2/3 Pathway *. *Journal of Biological Chemistry*, 276(28):26448–26452, July 2001. ISSN 0021-9258, 1083-351X. doi: 10.1074/jbc.M103856200. URL https://www.jbc.org/article/S0021-9258(19)30207-8/abstract. Publisher: Elsevier.

E. E. Sander, S. Van Delft, J. P. Ten Klooster, T. Reid, R. A. Van Der Kammen, F. Michiels, and J. G. Collard. Matrix-dependent Tiam1/Rac Signaling in Epithelial Cells Promotes Either Cell–Cell Adhesion or Cell Migration and Is Regulated by Phosphatidylinositol 3-Kinase. *The Journal of Cell Biology*, 143(5):1385–1398, Nov. 1998. ISSN 0021-9525, 1540-8140. doi: 10.1083/jcb.143.5.1385. URL https://rupress.org/jcb/article/143/5/1385/29443/Matrix-dependent-Tiam1-Rac-Signaling-in-Epithelial.

S. Srinivasan, F. Wang, S. Glavas, A. Ott, F. Hofmann, K. Aktories, D. Kalman, and H. R. Bourne. Rac and Cdc42 play distinct roles in regulating PI(3,4,5)P3 and polarity during neutrophil chemotaxis. *The Journal of Cell Biology*, 160(3):375–385, Feb. 2003. ISSN 1540-8140, 0021-9525. doi: 10.1083/jcb.200208179. URL https://rupress.org/jcb/article/160/3/375/54784/Rac-and-Cdc42-play-distinct-roles-in-regulating-PI.

Y. Sun, H. Do, J. Gao, R. Zhao, M. Zhao, and A. Mogilner. Keratocyte fragments and cells utilize competing pathways to move in opposite directions in an electric field. *Current Biology*, 23(7): 569–574, 2013. ISSN 09609822. doi: 10.1016/j.cub.2013.02.026. URL http://dx.doi.org/10.1016/j.cub.2013.02.026. ISBN: 3143627344 Publisher: Elsevier Ltd.

Y. H. Y. Sun, Y. H. Y. Sun, K. Zhu, B. Reid, X. Gao, B. W. Draper, M. Zhao, and A. Mogilner. Electric fields accelerate cell polarization and bypass myosin action in motility initiation. *Journal of Cellular Physiology*, 233(3):2378–2385, 2018. ISSN 10974652. doi: 10.1002/jcp.26109. ISBN: 9550161005.

T. M. Svitkina and G. G. Borisy. Arp2/3 Complex and Actin Depolymerizing Factor/Cofilin in Dendritic Organization and Treadmilling of Actin Filament Array in Lamellipodia. *The Journal of Cell Biology*, 145(5):1009–1026, May 1999. ISSN 0021-9525, 1540-8140. doi: 10.1083/jcb.145.5.1009. URL https://rupress.org/jcb/article/145/5/1009/16134/Arp2-3-Complex-and-Actin-Depolymerizing-Factor.

K. F. Tolias, J. H. Hartwig, H. Ishihara, Y. Shibasaki, L. C. Cantley, and C. L. Carpenter. Type I$\alpha$ phosphatidylinositol-4-phosphate 5-kinase mediates Rac-dependent actin assembly. *Current Biology*, 10(3):153–156, Feb. 2000. ISSN 09609822. doi: 10.1016/S0960-9822(00)00315-8. URL https://linkinghub.elsevier.com/retrieve/pii/S0960982200003158.

T. Tsuji, T. Ishizaki, M. Okamoto, C. Higashida, K. Kimura, T. Furuyashiki, Y. Arakawa, R. B. Birge, T. Nakamoto, H. Hirai, and S. Narumiya. ROCK and mDia1 antagonize in Rho-dependent Rac activation in Swiss 3T3 fibroblasts. *The Journal of Cell Biology*, 157(5):819–830, May 2002. ISSN 1540-8140, 0021-9525. doi: 10.1083/jcb.200112107. URL https://rupress.org/jcb/article/157/5/819/32739/ROCK-and-mDia1-antagonize-in-Rho-dependent-Rac.

F. Wang, P. Herzmark, O. D. Weiner, S. Srinivasan, G. Servant, and H. R. Bourne. Lipid products of PI(3)Ks maintain persistent cell polarity and directed motility in neutrophils. *Nature Cell Biology*, 4(7):513–518, July 2002. ISSN 1465-7392, 1476-4679. doi: 10.1038/ncb810. URL https://www.nature.com/articles/ncb810.

R. Wedlich-Soldner, S. Altschuler, L. Wu, and R. Li. Spontaneous Cell Polarization Through Actomyosin-Based Delivery of the Cdc42 GTPase. *Science*, 299(5610):1231–1235, Feb. 2003. doi: 10.1126/science.1080944. URL https://www-science-org.recursos.biblioteca.upc.edu/doi/full/10.1126/science.1080944. Publisher: American Association for the Advancement of Science.

O. D. Weiner, G. Servant, M. D. Welch, T. J. Mitchison, J. W. Sedat, and H. R. Bourne. Spatial control of actin polymerization during neutrophil chemotaxis. *Nature Cell Biology*, 1(2):75–81, June 1999. ISSN 1465-7392, 1476-4679. doi: 10.1038/10042. URL https://www.nature.com/articles/ncb0699_75.

O. D. Weiner, P. O. Neilsen, G. D. Prestwich, M. W. Kirschner, L. C. Cantley, and H. R. Bourne. A PtdInsP3- and Rho GTPase-mediated positive feedback loop regulates neutrophil polarity. *Nature Cell Biology*, 4(7):509–512, 2002. ISSN 14657392. doi: 10.1038/ncb811.

O. D. Weiner, W. A. Marganski, L. F. Wu, S. J. Altschuler, and M. W. Kirschner. An Actin-Based Wave Generator Organizes Cell Motility. *PLOS Biology*, 5(9):e221, Aug. 2007. doi: 10.1371/journal.pbio.0050221. URL https://doi.org/10.1371/journal.pbio.0050221. Publisher: Public Library of Science.

H. C. Welch, W. Coadwell, L. R. Stephens, and P. T. Hawkins. Phosphoinositide 3-kinase-dependent activation of Rac. *FEBS Letters*, 546(1):93–97, July 2003. ISSN 0014-5793, 1873-3468. doi: 10.1016/S0014-5793(03)00454-X. URL https://febs.onlinelibrary.wiley.com/doi/10.1016/S0014-5793%2803%2900454-X.

K. Wong, O. Pertz, K. Hahn, and H. Bourne. Neutrophil polarization: Spatiotemporal dynamics of RhoA activity support a self-organizing mechanism. *Proceedings of the National Academy of Sciences*, 103(10):3639–3644, Mar. 2006. ISSN 0027-8424, 1091-6490. doi: 10.1073/pnas.0600092103. URL https://pnas.org/doi/full/10.1073/pnas.0600092103.

R. A. Worthylake and K. Burridge. RhoA and ROCK Promote Migration by Limiting Membrane Protrusions. *Journal of Biological Chemistry*, 278(15):13578–13584, Apr. 2003. ISSN 00219258. doi: 10.1074/jbc.M211584200. URL https://linkinghub.elsevier.com/retrieve/pii/S0021925819647447.

J. Xu, F. Wang, A. Van Keymeulen, P. Herzmark, A. Straight, K. Kelly, Y. Takuwa, N. Sugimoto, T. Mitchison, and H. R. Bourne. Divergent signals and cytoskeletal assemblies regulate self-organizing polarity in neutrophils. *Cell*, 114(2):201–214, 2003. ISSN 00928674. doi: 10.1016/S0092-8674(03)00555-5.

T. Yeung, G. E. Gilbert, J. Shi, J. Silvius, A. Kapus, and S. Grinstein. Membrane phosphatidylserine regulates surface charge and protein localization. *Science (80-. ).*, 319(5860):210–213, 2008. ISSN 0036-8075. doi: 10.1126/science.1152066. URL http://science.sciencemag.org/content/319/5860/210.abstract. ISBN: 1095-9203 (Electronic).

B. Zhang and Y. Zheng. Regulation of RhoA GTP Hydrolysis by the GTPase-Activating Proteins p190, p50RhoGAP, Bcr, and 3BP-1. *Biochemistry*, 37(15):5249–5257, Apr. 1998. ISSN 0006-2960, 1520-4995. doi: 10.1021/bi9718447. URL https://pubs.acs.org/doi/10.1021/bi9718447.

M. Zhao, J. Pu, J. V. Forrester, and C. D. McCaig. Membrane lipids, EGF receptors, and intracellular signals colocalize and are polarized in epithelial cells moving directionally in a physiological electric field. *Faseb Journal*, 16(8):857–859, 2002. ISSN 15306860. doi: 10.1096/fj.01-0811fje.

M. Zhao, B. Song, J. Pu, T. Wada, B. Reid, G. Tai, F. Wang, A. Guo, P. Walczysko, Y. Gu, T. Sasaki, A. Suzuki, J. V. Forrester, H. R. Bourne, P. N. Devreotes, C. D. McCaig, and J. M. Penninger. Electrical signals control wound healing through phosphatidylinositol-3-OH kinase-$\gamma$ and PTEN. *Nature*, 442(7101):457–460, July 2006. ISSN 1476-4687. doi: 10.1038/nature04925. URL https://www.nature.com/articles/nature04925. Publisher: Nature Publishing Group.

Y. Zheng, S. Bagrodia, and R. A. Cerione. Activation of phosphoinositide 3-kinase activity by Cdc42Hs binding to p85. *Journal of Biological Chemistry*, 269(29):18727–18730, July 1994. ISSN 0021-9258. doi: 10.1016/S0021-9258(17)32226-3. URL https://www.sciencedirect.com/science/article/pii/S0021925817322263.

G. C. Zondag, E. E. Evers, J. P. Ten Klooster, L. Janssen, R. A. Van Der Kammen, and J. G. Collard. Oncogenic Ras Downregulates Rac Activity, Which Leads to Increased Rho Activity and Epithelial–Mesenchymal Transition. *The Journal of Cell Biology*, 149(4):775–782, May 2000. ISSN 0021-9525, 1540-8140. doi: 10.1083/jcb.149.4.775. URL https://rupress.org/jcb/article/149/4/775/43852/Oncogenic-Ras-Downregulates-Rac-Activity-Which.
